# Isoform diversity as a complementary dimension of gene expression

**DOI:** 10.64898/2026.11.05.764607

**Authors:** Shaked Shanas, Eliron Lubaton, Shaked Jermans, Martin Mikl

**Affiliations:** Department of Human Biology, School of Life Sciences, Faculty of Natural Sciences, University of Haifa, Haifa 3103301, Israel

## Abstract

Most human genes produce multiple mRNA isoforms through alternative splicing and alternative transcription start and end sites. Isoforms of the same gene can differ in function, and their relative abundance varies between tissues and conditions. However, gene expression analysis is often restricted to the gene level, masking its diversification into mRNA and protein isoforms with potentially distinct functionality. Here we show, using an adaptation of Shannon evenness applied to 4,814 GTEx samples from 25 tissues, that isoform diversity constitutes a distinct dimension of gene expression, complementary to gene-level abundance and whole transcriptome diversity. Isoform diversity separated tissues based on non-differentially expressed genes and changed with age in a gene- and tissue-specific manner, most strongly in the brain and often without changes in overall gene expression levels. During neuronal differentiation, it increased most notably in RNA processing and splicing genes. Our results suggest that isoform diversity represents an independent dimension of gene expression and may help explain functional differences between tissues and cell states that are not captured by analyzing expression levels alone.

## Introduction

Most human genes produce more than one transcript isoform, through alternative splicing and alternative start and end sites of transcription^1–3^. A gene’s output is therefore a distribution of isoforms, not a single quantity. The shape of this distribution can carry information about the diversification of gene function. In many cases, a single transcript is dominant^4,5^, but a substantial fraction of genes co-express several isoforms at comparable levels^6^. In general, the relative abundance of the isoforms can change across tissues and conditions^2,6,7^, often without changes in the overall mRNA expression level of the gene^8^. Isoforms of the same gene can differ markedly in function, localization and enzymatic activity^9^ and often share less than half of their interaction partners, resembling distinct proteins rather than variants of the same protein^10^. Tissue-regulated exons frequently remodel protein interactions^11^, often through binding motifs in disordered region^12^. The relative abundance of a gene’s isoforms can therefore serve as a proxy for its functional diversification, although not every detected isoform is translated or functional^13^.

Diversity of gene expression can be quantified at different levels. Transcriptome diversity describes how evenly expression is distributed across all genes of a sample^14,15^ and has been identified as a major hidden source of variation in RNA-seq data^16^. Isoform diversity describes the distribution of expression of a gene across its isoforms within a sample^15^. It can be summarized by the number of expressed isoforms, by variance-based measures such as the Fano factor, or by information-theoretic measures^15^. Shannon entropy^17^ was first used on microarray expression data to show that isoform diversity increases in cancer^18^. It has since been used for example to compare the isoform output of duplicated genes in the brain^19^ and to follow splicing changes in glioma^20^, and Jensen– Shannon divergence has been used to detect isoform switching during cell differentiation^7^. Entropy-based quantification of diversity has also been applied to individual alternative splicing events^8^ and different metrics have been compared for their applicability to gene expression data^21^. Perplexity^22^, the exponential of Shannon entropy, quantifies isoform diversity as an effective number of isoforms. Applying this approach to long-read data from the ENCODE4 project indicated that isoform diversity is largely uncoupled from gene expression and differs between gene groups^22^. These existing gene-level diversity metrics either depend on isoform number, such as perplexity^22^, or handle low coverage with fixed pseudo-counts or filtering^21^, with no single method having emerged as a standard^15,22^.

Here, we use a gene-level measure of isoform diversity (I-DIV), based on Shannon evenness with a dynamic pseudocount that accounts for differences in expression levels and isoform number, and apply it to the Genotype-Tissue Expression (GTEx) project and an independent neuronal differentiation dataset. We show that isoform diversity is a gene-specific property that varies with tissue identity, age and cell state largely independently of both expression level and transcriptome diversity.

## Results

### A gene-level measure of isoform diversity across human tissues

To describe how the expression of a genes is distributed across its annotated isoforms, we built a gene-level isoform diversity metric based on Shannon evenness, an entropy-based measure previously used to quantify gene expression diversity^14,18,23^. To account for the dependence of entropy-based diversity measures on sequencing depth and isoform number, we added a dynamic pseudo-count based on gene coverage (Fig. 1a, Methods). Importantly, we added this value to all possible isoforms that are annotated and theoretically could be expressed, not limiting it to the isoforms detected in a particular sample. This approach was chosen to account for effects of expression level and sampling depth on the number of detected isoforms (Methods). Simulating different sampling depths for the same underlying distribution of isoform expression of a gene (Supplementary Fig. 1a) showed that this isoform diversity metric (I-DIV) is largely independent of sampling depth (Supplementary Fig. 1b), allowing to estimate the “true” diversity from a low coverage sample. We applied this metric to transcript-level expression data from the Genotype-Tissue Expression project (GTEx v8) which provides RNA sequencing data from dozens of tissues in hundreds of donors (Fig. 1a). For each gene in each sample, we calculated its isoform diversity (I-DIV, Methods) and applied a number of gene- and sample-based filtering steps. Postmortem transcriptomes are sensitive to tissue quality and to the circumstances of death^24^. We therefore kept only samples with an RNA integrity number (RIN) above 6 and a Hardy score of 1 or 2 (fast violent or natural death, excluding ventilator cases and slow or intermediate death types), and only tissues represented by at least 15 samples. We restricted the analysis to protein-coding genes (based on GENCODE v26^25^) with a non-zero I-DIV in at least 1% of all samples (48), indicative of non-zero expression and the existence of at least two potential mRNA isoforms. The resulting dataset comprised 15,834 genes and 4,814 samples from 267 donors across 25 tissues (Fig. 1a), providing the scale and breadth to compare isoform diversity systematically across genes, tissues and individuals.

**Figure 1.**
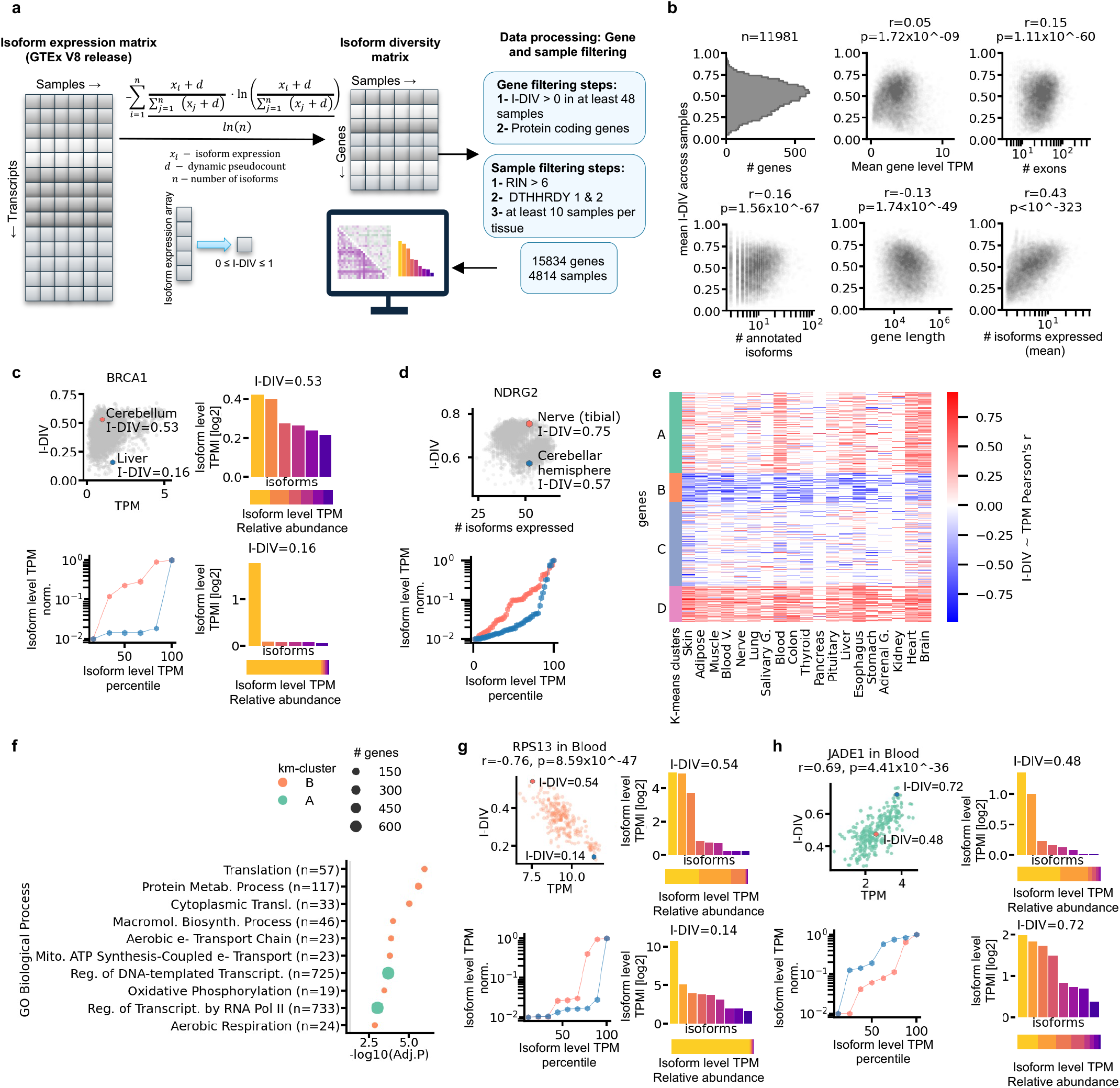
I-DIV summarizes the relative abundances of a gene’s isoforms and can be decoupled from gene expression. (a) Outline of data processing and filtering. The isoform expression matrix from GTEx V8 is aggregated per gene into an isoform diversity (I-DIV) matrix. (b) Overview of I-DIV in comparison with gene features. Every datapoint represents a different gene (n=11,981), mean measurements are across 4814 GTEx samples. Top panels, left to right: distribution of mean I-DIV; mean I-DIV plotted against mean gene expression(log2(TPM+1); TPM); mean I-DIV plotted against number of exons (# exons). Bottom panels, left to right: mean I-DIV plotted against mean number of annotated isoforms (# annotated isoforms), gene length, and mean number of expressed isoforms (# isoforms expressed). Pearson’s correlation coefficients and p-values are stated in each panel. (c) *BRCA1* as an example of I-DIV being decoupled from gene expression (log2(TPM+1)). Top left: I-DIV against gene expression across GTEx samples with I-DIV > 0 (n = 4,813; Pearson r = 0.3, p =1.21×10^−101^), with the Pearson correlation coefficient and two-sided p-value stated above. Highlighted hexagons mark a cerebellum sample colored in salmon (I-DIV=0.53; log2(TPM+1)= 0.97) and a liver sample in blue (I-DIV=0.16; log2(TPM+1)= 1.64), each expressing six isoforms. Top right: *BRCA1* isoform expression (log2(TPM+1)) in the cerebellum sample, with a stacked bar plot of relative isoform expression below. Bottom right: the same for the liver sample. Bottom left: isoform expression on scaled TPM; hexagons represent individual isoforms, and salmon and blue colored lines correspond to the two samples shown in the top-left and bottom-right panels. (d) *NDRG2* as an example of I-DIV being decoupled from the number of expressed isoforms. Top: I-DIV against the number of isoforms expressed, across samples with I-DIV > 0 and log2(TPM+1) > 5.5 (n = 4,161; r = −0.34, p=1.27×10^−115^). Highlighted hexagons mark a nerve sample in salmon color (I-DIV=0.75) and a cerebellar hemisphere sample in blue (I-DIV=0.57) that express the same number of isoforms (n = 52). Bottom: isoform expression on scaled TPM; hexagons represent individual isoforms, and orange and blue lines correspond to the two samples highlighted in the top panel. (e) Clustered heatmap of Pearson correlation coefficients between I-DIV and gene expression per somatic tissue (n genes = 15,834; n SMTS tissues = 19). Genes were grouped into four k-means clusters, A–D (n = 5,556, 1,975, 5,802 and 2,501 genes, respectively). (f) Enrichment analysis of each cluster using GSEApy Enrichr (n = 5,556, 1,975, 5,802 and 2,501 genes) against a background of all genes in the analysis (n = 15,834). Colors correspond to the k-means cluster bar in panel e. Results are shown for the top 10 GO Biological Process 2025 gene set terms on y axis, negative log10(FDR) as x axis; significantly enriched terms (FDR<0.05) shown from cluster A and B; the number of genes per term is expressed as the size. (g) *RPS13* as an example of a gene from the GO term Translation (GO:0006412) enriched in cluster B. Top left: I-DIV against gene expression across Blood samples with I-DIV > 0 (n = 225), with the Pearson correlation coefficient and two-sided p-value stated above. Highlighted hexagons mark samples with high I-DIV in salmon (I-DIV=0.54; log2(TPM+1)=7.54) and low I-DIV in blue (I-DIV=0.14; log2(TPM+1)=11.35) that express the same number of isoforms (n = 9). Right: isoform expression (log2(TPM+1)) in the sample with higher (top) or lower (bottom) I-DIV values, with a stacked bar plot of relative isoform expression below. Bottom left: isoform expression on scaled TPM; hexagons represent individual isoforms, and orange and blue lines correspond to the two samples at top left panel. (h) *JADE1* as an example of a gene from the GO terms Regulation of DNA-templated Transcription (GO:0006355) and Regulation of Transcription by RNA Polymerase II (GO:0006357) which are significantly enriched (FDR< 0.05) in cluster A. Top left: I-DIV against gene expression across blood samples with I-DIV > 0 (n = 245), with the Pearson correlation coefficient and two-sided p-value stated above. Highlighted hexagons mark samples with low I-DIV in salmon (I-DIV=0.48; log2(TPM+1)=2.58) and high I-DIV in blue (I-DIV=0.72; log2(TPM+1)=3.73) that express the same number of isoforms (n = 8). Right: isoform expression (log2(TPM+1)) in the sample with lower (top) or higher (bottom) I-DIV, with a stacked bar plot of relative isoform expression below. Bottom left: isoform expression on scaled TPM; hexagons represent individual isoforms, and orange and blue lines correspond to the two samples at top left panel.

### Isoform diversity is largely independent of gene expression and structure

To characterize the relationship of isoform diversity to other gene features, we compared mean I-DIV with expression level, gene structure and isoform number across 11,981 genes with on average more than two isoforms expressed (Fig. 1b). Mean I-DIV across 4,814 samples varied widely between genes and showed a unimodal and approximately symmetric distribution (median = 0.5, range 0.05– 0.95; Fig. 1b). I-DIV was almost independent of mean gene expression (Pearson’s r = 0.05) and was also only weakly associated with the number of exons (r = 0.15), the number of annotated isoforms (r = 0.16) and gene length (r = −0.13), with gene expression level showing comparable associations with these features (Supplementary Fig. 2a). I-DIV as well as gene-level TPM correlated more strongly with the mean number of expressed isoforms (r = 0.43 and r=0.4, resp.). However, also the number of expressed isoforms explained less than 20% of the variance in I-DIV (r² = 0.18), showing that the I-DIV metric is only partly determined by how many isoforms are expressed, but captures also the distribution of expression between these isoforms.

To investigate the relationship between I-DIV, gene expression level and number of isoforms in individual genes, we first focused on two well-known tumor suppressor genes as examples. In the case of BRCA1, expression and I-DIV were moderately correlated across samples and tissues (Fig. 1c; Pearson r = 0.3, p = 1.21×10^−101^), with a strong dependence of I-DIV on the number of isoforms expressed (Supplementary Fig. 2b, top). Comparing a liver and a cerebellum sample in which *BRCA1* was expressed at comparable levels (log2(TPM+1) = 1.64 and 0.97, respectively) and from six isoforms each, I-DIV differed substantially between the two samples (Fig. 1c). While in the liver sample a single isoform was dominant (I-DIV = 0.16), in the cerebellum sample *BRCA1* expression was more evenly distributed across several isoforms (I-DIV = 0.53). To visualize relative isoform abundances across samples and genes in a comparative manner, we plotted isoform-level TPMs for all isoforms ranked by their expression level and normalized to the expression of the most abundant isoform. This shows the difference in diversity as a steep profile in the liver (blue) and a flatter one in the cerebellum (salmon, Fig. 1c, bottom left). In the case of *NDRG2*, which expresses a particularly large number of isoforms, I-DIV was negatively correlated with expression levels (Supplementary Fig. 2b, bottom; Pearson r=-0.51, p = 6.17×10^−270^) and, counterintuitively, also with the number of isoforms expressed (Fig. 1d, top; Pearson r=-0.3, p = 1.27×10^−115^). A tibial nerve sample and a cerebellar hemisphere sample both expressed 52 isoforms of *NDRG2*, but they differed in I-DIV (0.75 and 0.57, respectively), reflecting a more even distribution of expression across isoforms in the nerve sample (salmon, Fig. 1d, bottom). Together, these results indicate that isoform diversity can be decoupled from expression level and gene architecture, both across genes and across samples of the same gene.

### The coupling between isoform diversity and expression is gene-specific and partly tissue-dependent

Across genes and tissues, mean I-DIV was almost unrelated to mean expression (Fig. 1b). To test whether this also holds within genes and tissues, we correlated I-DIV with gene expression across samples for each gene and tissue, and clustered genes by their correlation profiles (k-means; 15,834 genes, 19 tissues; Fig. 1e; Supplementary Data 1). Clusters B (1,975 genes) and D (2,501 genes) showed consistently negative and positive correlations across tissues, respectively. Clusters A (5,556 genes) and C (5,802 genes) showed weaker correlations that varied more between tissues. The clusters also differed in their mean I-DIV and in the strength of the correlation between number of expressed isoforms to I-DIV (Supplementary Fig. 2c). Only clusters A and B were enriched for functional terms (GSEApy Enrichr^26,27^; Fig. 1f and Supplementary Fig. 2c). Cluster A was enriched for genes involved in regulation of transcription and DNA repair, and cluster B for translation and aerobic respiration. In blood, for example, the ribosomal protein gene RPS13 (cluster B) showed negative correlation between I-DIV and expression level: a sample with higher expression had lower I-DIV and vice versa (log2(TPM+1) = 11.35, I-DIV = 0.14 versus log2(TPM+1) = 7.54, I-DIV = 0.54; Fig. 1g). Conversely, for the transcriptional regulator JADE1 (cluster A), higher expression was associated with more even isoform usage, e.g. in two samples both expressing eight isoforms (log2(TPM+1) = 3.73, I-DIV = 0.72 versus log2(TPM+1) = 2.58, I-DIV = 0.48; Fig. 1h). These results indicate that the relationship between isoform diversity and expression is a gene-specific property.

### Isoform diversity separates tissues based on equivalently expressed genes

As I-DIV is largely uncoupled from the overall gene expression level, we asked whether it carries additional tissue-specific information. For every pair of GTEx tissues (300 pairs of 25 tissues), we compared how well principal component analyses (PCAs) separated the two tissues based on either gene expression (log2(TPM+1)) or I-DIV, using silhouette scores as a metric for the separation between samples from the two tissues (Fig. 2a, Methods). To minimize the contribution of differences in gene expression levels, we repeated each comparison using only genes with no evidence for differential expression between the two tissues (nonDE; equivalence testing using two one-sided tests on limma^28^ estimates, margin ±0.5 log2(TPM+1), FDR < 0.05; Methods). Using all 15,834 genes, both expression and I-DIV separated most tissue pairs well (Fig. 2b), with TPM-based separation performing slightly better than I-DIV-based separation (Fig. 2c). Restricting the analysis to nonDE genes largely abolished separation by expression. I-DIV profiles of the same genes, however, still allowed separation for many tissue pairs (Fig. 2bc; median silhouette score 0.02 (TPM) vs 0.4 (I-DIV) on nonDE genes, p=3.48×10^−45^, Wilcoxon test).

**Figure 2.**
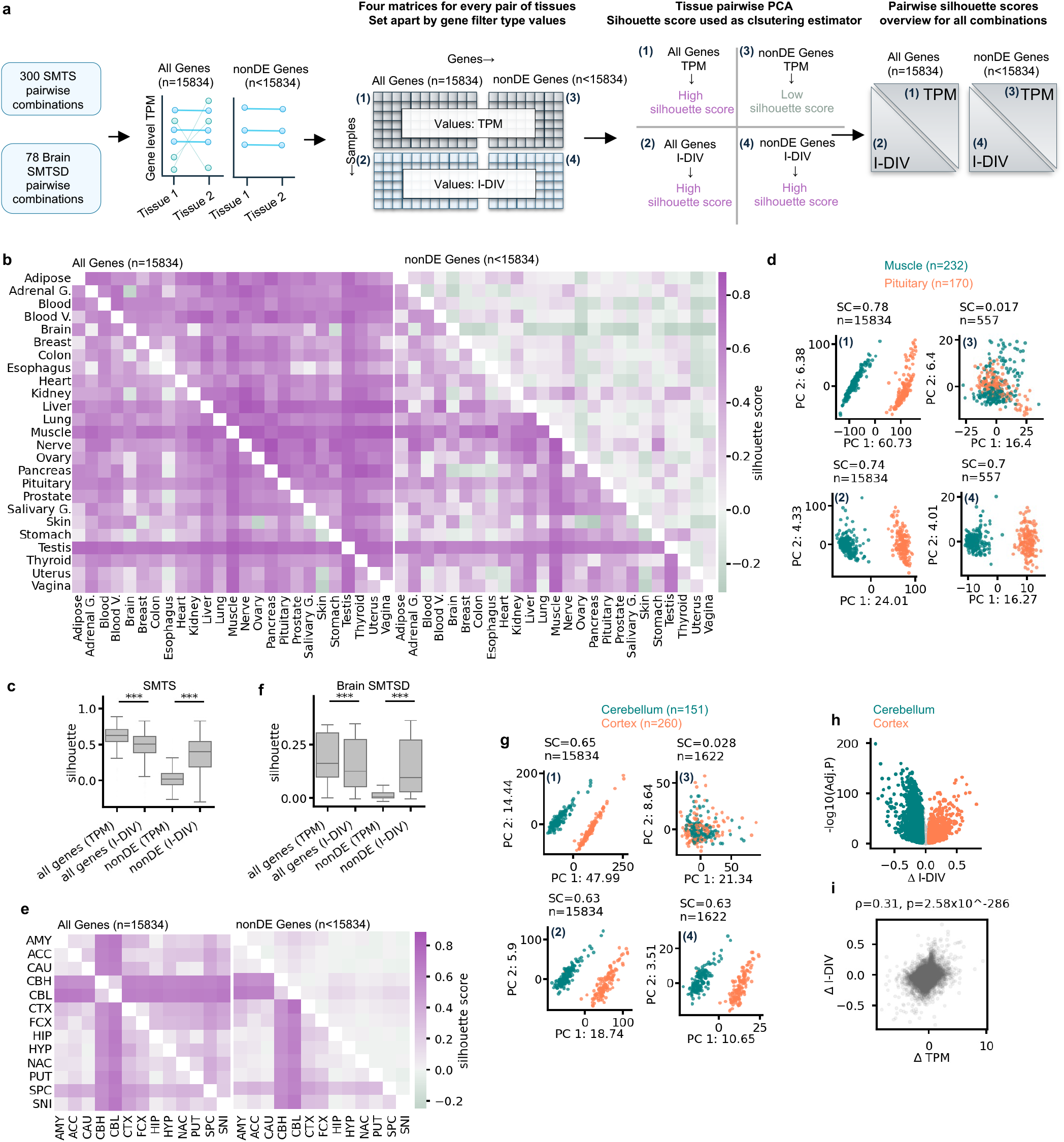
Isoform diversity separates tissues based on non-differentially expressed genes. (a) Overview of the pairwise tissue comparison. For each pair, genes were assigned to two groups: all genes in the analysis (*n* = 15,834), and genes with statistically equivalent expression between the two tissues (nonDE genes; *n* < 15,834, pair-specific). Four sample-by-gene matrices per pair were generated: (1) gene expression (TPM) of all genes, (2) I-DIV of all genes, (3) gene expression of nonDE genes and (4) I-DIV of nonDE genes. Each matrix was reduced by PCA (number of components=2), and separation of the two tissues was scored with the silhouette coefficient. (b) Overview of pairwise PCAs of SMTS tissues (number of components=2), scored by silhouette. Left heatmap: PCAs based on all genes in the analysis (n = 15,834). Right heatmap: PCAs based on nonDE genes (different for each pair). Values above the diagonal show scores for PCAs based on gene expression (TPM); values below the diagonal show scores for PCAs based on I-DIV. Number of SMTS tissues = 25; number of samples = 4,814; silhouette scores range from −0.3 to 0.89. (c) Silhouette scores of SMTS pairs as shown in (b). Box plots show the median (central line) and interquartile range (box, 25^th^–75^th^ percentiles), with whiskers extending to the most extreme data point within 1.5x of the interquartile range. Wilcoxon signed-Rank tests between silhouette scores of I-DIV against gene level TPM based on same gene filtering criteria (all genes p-value=1.12×10^−39^; nonDE p-value=3.48×10^−45^). (d) Pairwise PCA and silhouette scores for muscle (green; n = 232 samples) and pituitary gland samples (orange; n = 170 samples). Top left (1): PCA of gene expression across all protein-coding genes (n = 15,834; silhouette score = 0.78). Bottom left (2): PCA of I-DIV across all protein-coding genes (n = 15,834; silhouette score = 0.74). Top right (3): PCA of gene expression of non-DE genes (n = 557; silhouette score = 0.0.017). Bottom right: PCA of I-DIV of the same genes (n = 557; silhouette score = 0.70). (e) Overview of pairwise PCAs of SMTSD brain tissues (number of components=2), scored by silhouette. Left heatmap: PCAs based on all genes in the analysis (n = 15,834). Right heatmap: PCAs based on genes with statistically equivalent expression between the two tissues of each pair. Values above the diagonal show scores for PCAs based on gene expression; values below the diagonal show scores for PCAs based on I-DIV. Number of SMTSD brain tissues = 13; number of samples = 1,454; silhouette scores range from −0.03 to 0.69. Brain SMTSD tissues abbreviations: ACC, anterior cingulate cortex (BA24); AMY, amygdala; CAU, caudate (basal ganglia); CBH, cerebellar hemisphere; CBL, cerebellum; CTX, cortex; FCX, frontal cortex (BA9); HIP, hippocampus; HYP, hypothalamus; NAC, nucleus accumbens (basal ganglia); PUT, putamen (basal ganglia); SNI, substantia nigra; SPC, spinal cord (cervical c-1). (f) Silhouette scores of SMTSD pairs as shown in (e). Box plots show the median (central line) and interquartile range (box, 25^th^–75^th^ percentiles), with whiskers extending to the most extreme data point within 1.5x the interquartile range. Wilcoxon signed-Rank tests between silhouette scores of I-DIV and gene level TPM based on same gene filtering criteria (all genes p-value=6.47×10^−10^; nonDE p-value=1.68×10^−14^). (g) Pairwise PCA and silhouette scores for cerebellum (teal colored; n = 151 samples) and cortex (coral colored; n = 260 samples). Top left: PCA of gene expression across all protein-coding genes (n = 15,834; silhouette score = 0.65). Top right: PCA of I-DIV across all protein-coding genes (n = 15,834; silhouette score = 0.63). Bottom left: PCA of gene expression of genes with statistically equivalent expression between the two tissues (n = 1,622; silhouette score = 0.028). Bottom right: PCA of I-DIV of the same genes (n = 1,622; silhouette score = 0.63). (h) Volcano plot of the change in I-DIV (cortex-cerebellum) against the p-value corrected for multiple testing (tested using limma, FDR, Benjamini– Hochberg). Genes with median I-DIV above 0 across cortex and cerebellum samples were used (n = 13,328). Cut-offs for significance were adjusted p ≤ 0.05 and an absolute difference ≥ 0.05. Teal colored: genes with higher I-DIV in cerebellum (n = 5,115); Coral colored: genes with higher I-DIV in cortex (n = 1,853). (i) Difference in I-DIV (Δ I-DIV) against difference in log2(TPM+1) (Δ TPM) between cerebellum and cortex, for genes with median I-DIV>0 (n = 13,328). Spearman’s correlation coefficient and p-value are stated above.

One representative example is the distinction between muscle and pituitary gland samples, where the silhouette score decreased from 0.78 to 0.017 for TPM but only modestly from 0.74 to 0.7 for I-DIV when only the 557 nonDE genes were used (Fig. 2d). We then focused on more closely related tissues and tested whether I-DIV could also separate between brain regions (78 pairs of 13 regions; Fig. 2e). Here the ability to separate between sub-tissues was region-specific: Based on all genes, the distinction between cerebellar and non-cerebellar samples showed the highest silhouette scores for both TPM and I-DIV (Fig. 2e), which was maintained when using I-DIV, but not TPM values of nonDE genes (Fig. 2ef; Supplementary Data 2). Cerebellum and cortex, for example, were barely separated by the expression of their 1,622 nonDE genes (silhouette 0.028) but were clearly separated based on the I-DIV values of the same genes (0.63; Fig. 2g). These results show that genes expressed at the same level in two tissues can still differ systematically in how that expression is distributed across isoforms, in line with the extensive tissue dependence of isoform usage^2^, and that I-DIV can identify these systematic differences between tissues.

### Isoform diversity and expression differ in distinct gene sets between cerebellum and cortex

To characterize the differences at the gene level, we compared I-DIV between cerebellum and cortex using limma^28^ (13,328 genes with median I-DIV>0). More than half of the tested genes differed significantly in I-DIV (6,968 genes; 5,115 higher in cerebellum and 1,853 higher in cortex; FDR<0.05; Fig. 2h). Changes in I-DIV were only moderately correlated with changes in expression (Spearman’s ρ = 0.31; Fig. 2i). Differentially expressed genes and genes with significantly different I-DIV were mostly associated with distinct biological processes (Supplementary Fig. 3). Genes with higher I-DIV in cerebellum were enriched for protein catabolism, including ER-associated degradation and proteasomal processes. Genes with higher I-DIV in cortex were enriched for cilium movement, inflammatory response and GPCR signaling. Genes with higher expression were instead enriched for transcriptional regulation and chromatin organization in cerebellum, and for synaptic transmission and neuronal differentiation in cortex. Terms enriched for I-DIV and for expression overlapped only partly in cortex (10 shared terms) and not at all in cerebellum (Supplementary Fig. 3b). Isoform diversity can therefore reveal differences in gene expression between brain regions that are largely separate from those captured by differential expression analyses.

### Transcriptome diversity varies between tissues and partly reflects cellular composition

To compare the diversification of gene expression on the gene and the transcriptome level, we applied the same metric used for calculation of I-DIV to describing the diversity of transcriptomes. For each sample, we calculated transcriptome diversity (T-DIV) as the evenness of gene expression across all protein-coding genes, using the same approach of adding a dynamic pseudo-count to each gene (Fig. 3a; Methods). Related entropy-based measures of transcriptome diversity have been used to describe tissue specialization^14^ and have been identified as a major source of variation in RNA-seq data^16^. In the GTEx samples analyzed, T-DIV ranged from 0.17 in a blood sample to 0.91 in a testis sample (Fig. 3b). T-DIV differed between tissues (one-way ANOVA, p < 10^−323^; Fig. 3c). It was lowest in pancreas, indicative of the dominance of a group of highly expressed transcripts, and highest in testis, indicative of more even expression across genes, in accordance with earlier findings based on a different cohort and method^14^. This result was robust to exclusion (Fig. 3c) or inclusion (Supplementary Fig. 4a) of genes encoding ribosomal proteins, which, as a group, are among the most highly and broadly expressed genes across tissues and could dominate transcriptome-wide diversity measures.

**Figure 3.**
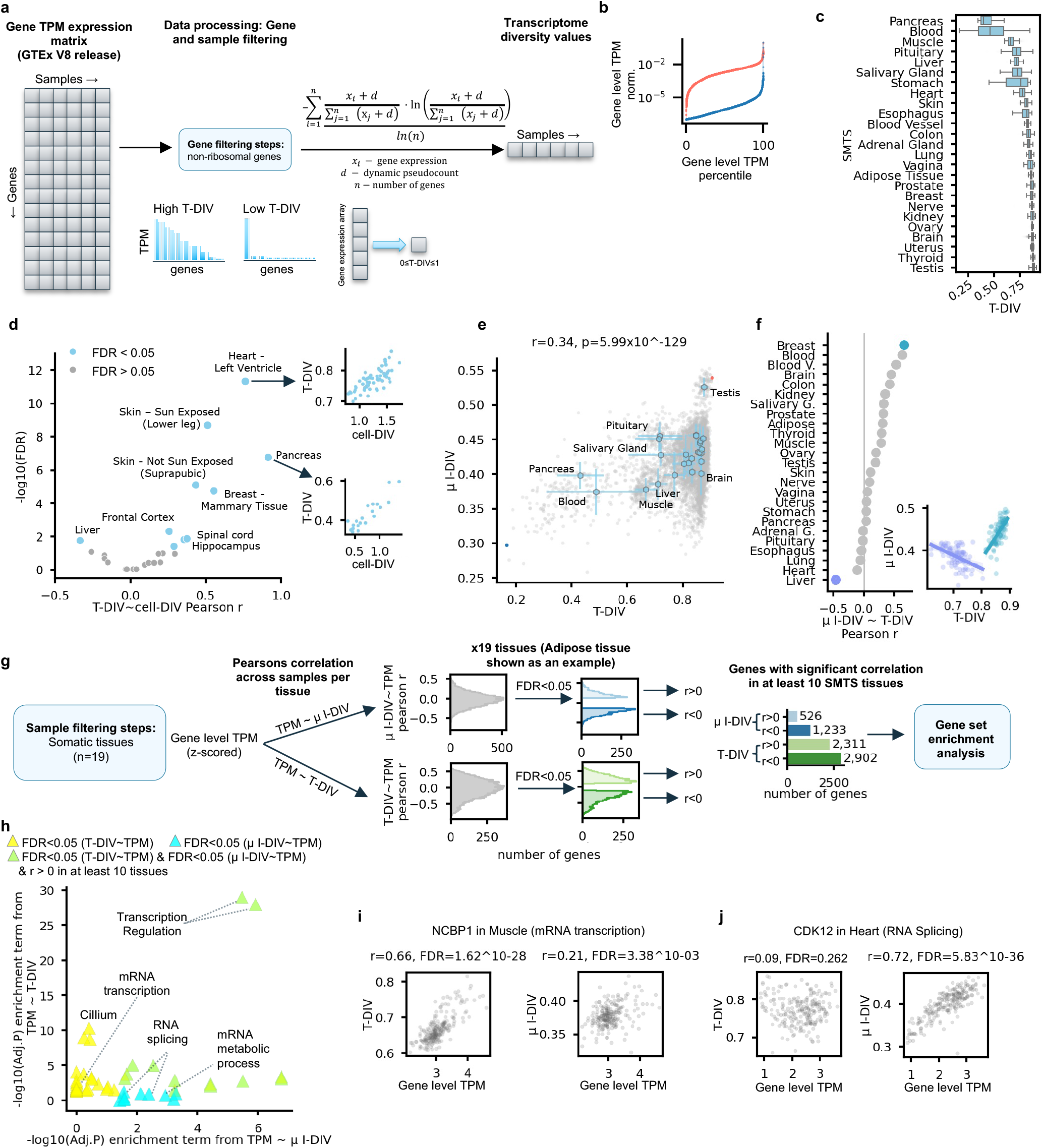
Isoform diversity is decoupled from transcriptome diversity and associated with a distinct set of regulators. (a) Outline of data processing. The gene expression matrix is first filtered to remove ribosomal genes and then aggregated into a single value per sample, the transcriptome diversity (T-DIV), which reflects the gene expression profile across the transcriptome calculated using the same formula for isoform diversity (Methods). (b) Lineplot of scaled gene expression ordered by expression TPM in samples with the highest and lowest T-DIV in the GTEx dataset. Every line represents a different sample. The sample with the highest T-DIV (salmon) originates from testis (SMTS) and has a value of 0.91, with n = 15,765 genes with TPM > 0. The sample with the lowest T-DIV (blue) originates from blood (SMTS) and has a value of 0.17, with n = 13,527 genes with TPM > 0. (c) T-DIV values across 25 SMTS tissues (T-DIV calculated from 15,690 genes. Box plots show the median (central line) and interquartile range (box, 25^th^–75^th^ percentiles), with whiskers extending to the most extreme data point within 1.5x of the interquartile range. One-way ANOVA: p < 10^−323^. (d) Pearson correlation of T-DIV with estimated cell diversity. Left: volcano plot showing Pearson correlation coefficients compared to corrected p-values (FDR, Benjamini–Hochberg). Each datapoint represents an SMTSD tissue; light blue points indicate SMTSD with significant correlations (FDR < 0.05) and grey points indicate non-significant correlations (FDR ≥ 0.05). Right: T-DIV against estimated cell diversity (cell-DIV) for heart (left ventricle) samples (top; n= 88; Pearson r= 0.76, FDR = 4.56×10-12) and pancreas samples (bottom; n= 26; Pearson r= 0.91, FDR = 1.67×10-07). (e) Mean I-DIV (µ I-DIV) against T-DIV per sample. Light grey points represent individual GTEx samples (n = 4,814); light blue hexagons represent the mean µ I-DIV and mean T-DIV per SMTS tissue, and light blue perpendicular lines show the standard deviation along each axis. Pearson’s correlation coefficient and two-sided p-value are stated above. (f) Left: Pearson correlation coefficient between µ I-DIV and T-DIV for each SMTS tissue; liver and breast, colored in violet and turquoise, have the strongest negative and positive correlations, respectively. Right: µ I-DIV against T-DIV for liver (n = 88; r= −0.46; p= 7×10^−06^) and breast (n = 93;r=0.65,p= 3.68×10^−13^) samples only, with regression lines per tissue; point colors match the highlighted tissues in the left panel. (g) Schematic of the analysis correlating gene expression with µ I-DIV or T-DIV per SMTS tissue (15,834 genes; 19 tissues). Genes were grouped by significant positive or negative correlation with µ I-DIV or with T-DIV in at least 10 of the 19 tissues: positive and negative correlation with µ I-DIV (light and dark blue, respectively) and positive and negative correlation with T-DIV (light and dark green, respectively; FDR <0.05). Each group underwent enrichment analysis using against a background of all 15,834 genes (GSEApy enrichr tool; FDR < 0.05). (h) Enrichr adjusted p-values of terms resulting from positively correlating genes with µ I-DIV and/or T-DIV. −log10(FDR) of the enrichment of GO terms among genes correlated with T-DIV (y axis) are plotted against −log10(FDR) of the enrichment of GO terms among genes correlated with µ I-DIV (x axis). Each point represents one significantly enriched term (Enrichr FDR < 0.05) in at least one of the axes. Blue points represent terms significantly enriched among genes correlating with µ I-DIV only; yellow points represent terms significantly enriched among genes correlating with T-DIV only; green points represent terms significantly enriched among genes correlating with both metrics. (i) T-DIV and µ I-DIV plotted against *NCBP1* (GO: mRNA transcription) expression in muscle (n = 232; T-DIV left, µ I-DIV right). Pearson’s correlation coefficients and two-sided p-values are stated above. (j) T-DIV and µ I-DIV plotted against *CDK12* (GO: RNA splicing) expression in heart (n = 225; T-DIV left, µ I-DIV right). Pearson’s correlation coefficients and two-sided p-values are stated above.

Because tissues differ in their cellular composition, we asked whether T-DIV reflects the diversity of cell types in a sample, using cell-type proportions estimated for GTEx samples by deconvolution^29^. T-DIV correlated positively with the estimated cell-type diversity (cell-DIV, based on Shannon entropy, Methods) in a subset of tissues, most strongly in pancreas and heart left ventricle, but not in others such as liver and frontal cortex (Fig. 3d). T-DIV therefore captures a sample-level property of the transcriptome that is tissue-specific and, in some cases, partly explained by cellular heterogeneity.

### The expression level of genes involved in RNA processing is positively correlated with mean isoform diversity

Because transcriptome diversity is a systematic source of variation across samples^16^, we tested whether it accounts for variation in isoform diversity, summarized as the mean I-DIV across genes (µ I-DIV). Across all samples, µ I-DIV and T-DIV were moderately correlated (Pearson’s r = 0.34, P = 5.99×10^−1^^29^; Fig. 3e), which was largely driven by tissues with low T-DIV such as blood and pancreas that also had low µ I-DIV. Within tissues, however, the relationship between mean I-DIV and T-DIV varied widely, showing positive correlation in breast (r = 0.65, p = 3.68×10^−13^) and negative correlation in liver (r = −0.46, p = 7×10^−6^; Fig. 3f). To identify genes whose expression level is robustly associated with either metric, we correlated the expression (log2(TPM+1)) of individual genes with T-DIV or µ I-DIV across samples in each tissue and selected genes that showed a significant correlation in at least 10 out of 19 tissues (FDR < 0.05; Fig. 3g). More genes were consistently associated with T-DIV (2,311 positive, 2,902 negative) than with µ I-DIV (526 positive, 1,233 negative). These four groups were enriched for partly distinct functional categories (Fig. 3h, Supplementary Fig. 4b, Supplementary Data 3), with no GO term being positively associated with one and negatively associated with the other metric. While terms related to oxidative phosphorylation were enriched in genes negatively correlated with both metrics (Supplementary Fig. 4b), transcriptional regulation GO terms were enriched in genes positively correlated with both metrics (Fig. 3h). GO terms specifically enriched among genes whose expression is positively correlated with T-DIV included mRNA transcription (Supplementary Fig. 4c). For example, expression of the cap binding protein NCBP1 (regulation) in muscle correlated strongly with T-DIV (r = 0.66, FDR = 1.62×10^−28^) and more weakly with µ I-DIV (r = 0.21, FDR = 3.38×10^−3^; Fig. 3i). In addition, mRNA splicing and metabolism annotations were specifically enriched in genes whose expression positively correlated with µ I-DIV (Fig. 3h, Supplementary Fig. 4d). For example, expression of the splicing-associated kinase CDK12 in heart correlated strongly with µ I-DIV (r = 0.72, FDR = 5.83×10^−36^) but not with T-DIV (r = 0.09, FDR = 0.262; Fig. 3j). Together, these results indicate that isoform diversity is associated with a distinct set of regulators, including components of the splicing machinery.

### Isoform diversity changes with age in a gene-and tissue-specific manner

Age-related changes have mostly been described based on gene-level expression^30–32^ or alternative splice isoform ratios^33–35^ or as increasing heterogeneity between individuals^36–38^. Using our gene- and transcriptome-level diversity metrics we asked whether the diversity of gene expression changes with age, globally or in a gene-specific manner. Because GTEx contains relatively few young donors, we combined age decades into three groups of more balanced size (20–49, 50–59 and 60–79 years; Fig. 4a; Methods). Mean I-DIV and T-DIV showed significant changes with age in only some tissues, and the direction of change often differed between the two measures (Fig. 4b). This agrees with the previously reported organ-specific character of transcriptomic changes with age^31,39,40^. In muscle, for example, both mean I-DIV and T-DIV increased with age (Spearman’s correlations: rho=0.21, p=0.001 and rho=0.15, p=0.02, resp.), while in brain, mean I-DIV decreased with age (Spearman’s correlations: rho= −0.11, p= 1.55×10^−05^) with no signficant changes in T-DIV (Fig. 4c and Supplementary Fig. 5a). When analyzed brain regions separately, a significant age-associated decrease in mean I-DIV was observable in Substantia nigra, Nucleus accumbens, Frontal Cortex and Anterior cingulate cortex, with similar trends in most other brain regions (Supplementary Fig. 5b). Taken together, changes in isoform diversity with age can be decoupled from changes in overall transcriptome diversity.

**Figure 4.**
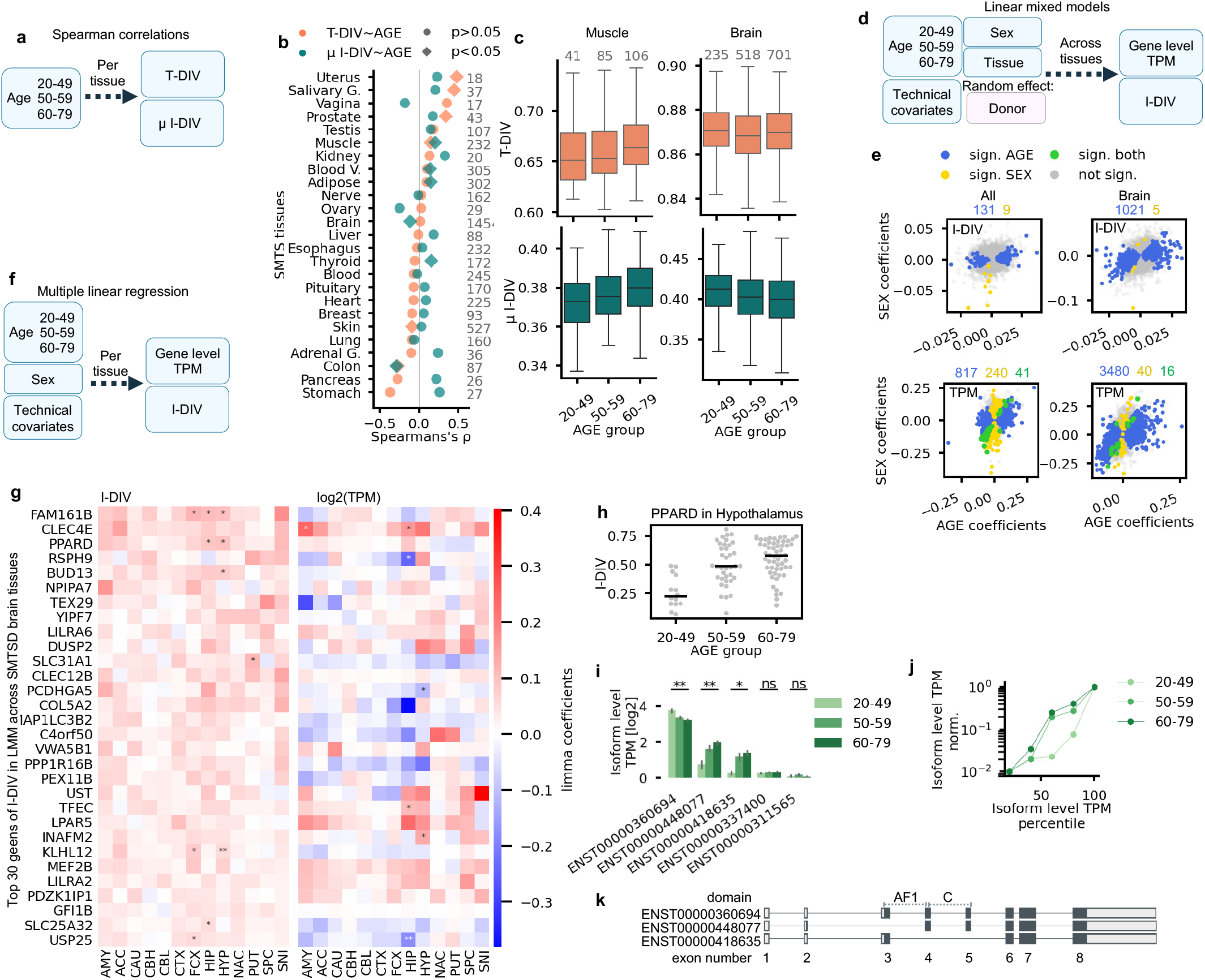
Isoform diversity changes with age in a tissue-and gene-specific manner. (a) Outline of Spearman correlation analysis between age and transcriptome diversity (T-DIV) or mean isoform diversity (µ I-DIV) for every tissue. (b) Spearman’s correlation coefficients of µ I-DIV (green) or T-DIV (orange) with age for each SMTS tissue. Diamonds indicate significant coefficients (two-sided p-value < 0.05) and circles indicate non-significant coefficients (two-sided p-value ≥ 0.05). The number of samples per tissue is stated in grey. (c) Box plots of the distribution of T-DIV (top) and µ I-DIV (bottom) in different age groups for muscle (left) and brain (right), showing the median (central line) and interquartile range (box, 25^th^–75^th^ percentiles), with whiskers extending to the most extreme data point within 1.5x of the interquartile range. Number of samples per age group is stated above each box. (d) Outline of linear mixed models applied to either gene level TPM or I-DIV using fixed effects of age (grouped to three groups), sex, tissue and technical covariates. Donor was included as random effect. (e) Comparison of age and sex coefficients of linear mixed models for I-DIV (top) and TPM (bottom); left: across all samples, with SMTS as fixed effect; right: across brain samples with SMTSD as fixed effect. Blue points: genes with a significant AGE coefficient (FDR < 0.05); yellow points: genes with a significant SEX coefficient (FDR < 0.05); green points: genes with significant coefficients for both AGE and SEX. Grey points: genes with no significant AGE nor SEX coefficient (FDR ≥ 0.05); number of genes are stated above each plot, with the color corresponding to the groups described above. (f) Outline of multiple linear regression on either gene level TPM or I-DIV, calculated per tissue. Covariates used: Age (grouped to three), sex and technical covariates. (g) Heatmaps show the age coefficients from multiple linear regression per gene and brain SMTSD tissue in I-DIV (left) and gene expression log(TPM+1) (right) for the top 30 genes ranked by I-DIV age coefficient which showed significance in I-DIV but not in TPM (FDR < 0.05) from the linear mixed model applied to brain regions. Asterisks indicate significant effects (Benjamini–Hochberg): ** FDR < 0.01; * FDR < 0.05. Brain SMTSD tissues: ACC, anterior cingulate cortex (BA24); AMY, amygdala; CAU, caudate (basal ganglia); CBH, cerebellar hemisphere; CBL, cerebellum; CTX, cortex; FCX, frontal cortex (BA9); HIP, hippocampus; HYP, hypothalamus; NAC, nucleus accumbens (basal ganglia); PUT, putamen (basal ganglia); SNI, substantia nigra; SPC, spinal cord (cervical c-1). (h) I-DIV of *PPARD* in hypothalamus samples, grouped by age brackets. Number of samples per age group, in x-axis order: 14, 37 and 56. (i) Bar plot of *PPARD* isoform mean expression per age group in the hypothalamus. Error bars show the standard error; age groups are colored in shades of green. Significance of isoform changes measured in p-values from multiple linear regression (Methods): ** p < 0.01; * p < 0.05; ns, not significant. (j) Line plot of the scaled mean expression of the *PPARD* isoforms, ordered by percentile of isoform expression across samples, for each age group in the hypothalamus (n=14, 37 and 56); age groups are colored in shades of green. (k) Exon (boxes) and intron (lines) structure of *PPARD* transcripts, adapted from the illustration available at the GTEx Portal. Transcripts: ENST00000360694, ENST00000448077, ENST00000418635. Dark boxes indicate coding sequences, domain names and approximate locations are shown above, exon numbers below.

To identify genes whose isoform diversity changes with age, we fitted linear mixed models of I-DIV and expression levels (TPM) for each gene. The models included tissue, sex, age, RIN and ischemic time as fixed effects and donor as a random effect, following previous approaches^32^, while adding technical covariates and tissue-level quantile normalization (Fig. 4d, Methods, Supplementary Data 4). Age effects on gene expression were highly consistent with those reported previously for GTEx^32^ (r = 0.94, p=2.63×10^−197^ for 424 genes significant in both analyses; Supplementary Fig. 5c, top), also when including the additional technical covariates (r = 0.92, p=1.39×10^−70^ for 176 genes significant in both analyses; Supplementary Fig. 5c, bottom). Changes in I-DIV of 131 genes were significantly associated with age (FDR < 0.05; Fig. 4e, left panels). Their age coefficients were only weakly correlated to the age coefficients for expression (r = 0.19; Supplementary Fig. 5d). Sex, in contrast, was associated with I-DIV of only 9 genes, compared with the expression level of 240 genes (Fig. 4e, left). Sex differences in expression therefore extend only rarely to isoform diversity.

Since brain showed I-DIV-specific age related effects across many of its 13 sampled regions (Supplementary Fig. 5b), we next tested for coordinated age-related changes across the brain. We applied the same LMM-based approach to identify significant contributions of age and sex to gene expression and isoform diversity variation (Fig. 4d, Supplementary Fig. 5e). The number of significant genes was higher than in the whole-body approach (Fig. 4e, right vs. left panels), arguably because age effects are more consistent across brain regions than across tissues. Like in the cross-tissue model, age dominated over sex with only 5 genes showing sex-associated changes in I-DIV compared to 1021 genes for age.

Because linear mixed models estimate one age effect per gene across tissues or brain regions, we next resolved age-related changes within individual subtissues (SMTSD) by multiple linear regression using limma^28^ and the same covariates as in the LMM-approach across tissues (Fig. 4f, Methods formula 6). In addition, we validated the results with a naïve approach, using Spearman correlation for I-DIV or log2(TPM+1) across all three age groups (Supplementary Fig 5f, Methods). Age-associated changes in I-DIV were concentrated in brain regions, which accounted for 87.32% (668 of 765) significant associations for I-DIV but only 55.41% (4259 of 7687) for expression (limma; FDR < 0.05; Supplementary Data 5). This is consistent with the extensive age-related changes in expression and splicing reported for the human brain^38,41,42^, and indicates that isoform diversity is particularly likely to change with age in the brain.

### Increasing PPARD isoform diversity with age favors isoforms with altered N-terminal domains

We next examined the genes with the strongest positive age effects on I-DIV in the brain-wide linear mixed models, restricted to those with no significant age effect on expression (I-DIV FDR < 0.05, TPM FDR ≥ 0.05). In both the multiple linear regression and Spearman analyses on individual brain regions, the I-DIV of these genes increased across most brain regions, although the increase reached significance (FDR < 0.05) in only some cases. Their expression, in contrast, typically did not change significantly with age and showed inconsistent trends across brain regions (Fig. 4g; Spearman’s correlation in Supplementary Fig. 5f).

At the top of the list was *FAM1C1B*, which encodes a centrosomal and ciliary protein. In GTEx v8, it has two mRNA isoforms and showed a consistent increase in the minor isoform at the expense of the major one across brain regions (Supplementary Fig. 5g). As an example of a gene with more isoforms, we examined *PPARD* whose I-DIV in the hypothalamus increased with age (coefficient = 0.1, FDR = 0.02; median I-DIV per age group: 20-49: 0.26, 50-59: 0.50, 60-79: 0.56; Fig. 4h). *PPARD* encodes PPARβ/δ, a nuclear receptor that is widely expressed in the brain and regulates lipid and energy metabolism, myelination and neuroinflammation^43,44^. The age-associated increase in I-DIV was a result of the dominant isoform decreasing and two alternative isoforms increasing in expression with age (Fig. 4ij). Both minor isoforms alter the N terminus of the encoded protein (Fig. 4k). ENST00000418635 lacks exons 4 and 5, containing the C domain consisting of two zinc-fingers through which PPARs bind their DNA response elements^45^. The other minor isoform, ENST00000448077, lacks exon 3 and differs in the translation start site (on exon 2 instead of 3), leading to an N terminus different from the canonical AF1 domain, an intrinsically disordered region responsible for ligand-independent activation^46^. *PPARD* isoforms have previously been shown to differ in their translation and activity^47^. The age-related increase in *PPARD* isoform diversity reflects a shift towards N-terminal variants with altered functionality and consequently may lead to changes in overall PPARδ activity. Among PPARs, this trend was specific to *PPARD* (Supplementary Fig. 5h) and occurred in most brain regions but not consistently outside the brain (Supplementary Fig. 5i). Together, these examples show that age-related changes in gene-level isoform diversity can point to specific shifts in isoform expression that remove or replace functional protein domains, even when the overall expression level of the gene remains unchanged.

### The isoform diversity of RNA processing genes increases during neuronal differentiation

To apply our analytical framework to an independent dataset, and to test whether regulatory patterns observed in human tissue samples also hold during a controlled cell-state transition, we analyzed a time course of human iPSC-derived neuronal differentiation^48^. RNA-seq and mass spectrometry data covered days 0–5 (4–5 replicates per day) from the start of transcription factor-induced differentiation^49^ (Fig. 5a). T-DIV increased steadily over differentiation (Spearman’s ρ = 0.98, p = 5.64×10^−17^; Fig. 5b), with mean I-DIV, mean number of isoforms per gene and number of genes expressed showing different trends over time (Supplementary Fig. 6a). Mean I-DIV increased over the first 24 hours and decreased again towards the end of the differentiation time course, which corresponds to the times of the largest morphological shifts where also the largest number of isoform changes could be detected^48^. The increase in T-DIV was also present when ribosomal protein genes were included (Supplementary Fig. 6b), indicating that it is not caused merely by a declining dominance of ribosomal protein transcripts. We then calculated a proteome diversity metric (P-DIV) from protein abundances as measured by mass spectrometry (Methods) and observed that P-DIV was strongly correlated with T-DIV (Spearman’s ρ = 0.9; p = 0.037; Fig. 5c). Transcriptome diversity is therefore largely reflected in the diversity of the proteome, with additional regulation at the translational and posttranslational level.

**Figure 5.**
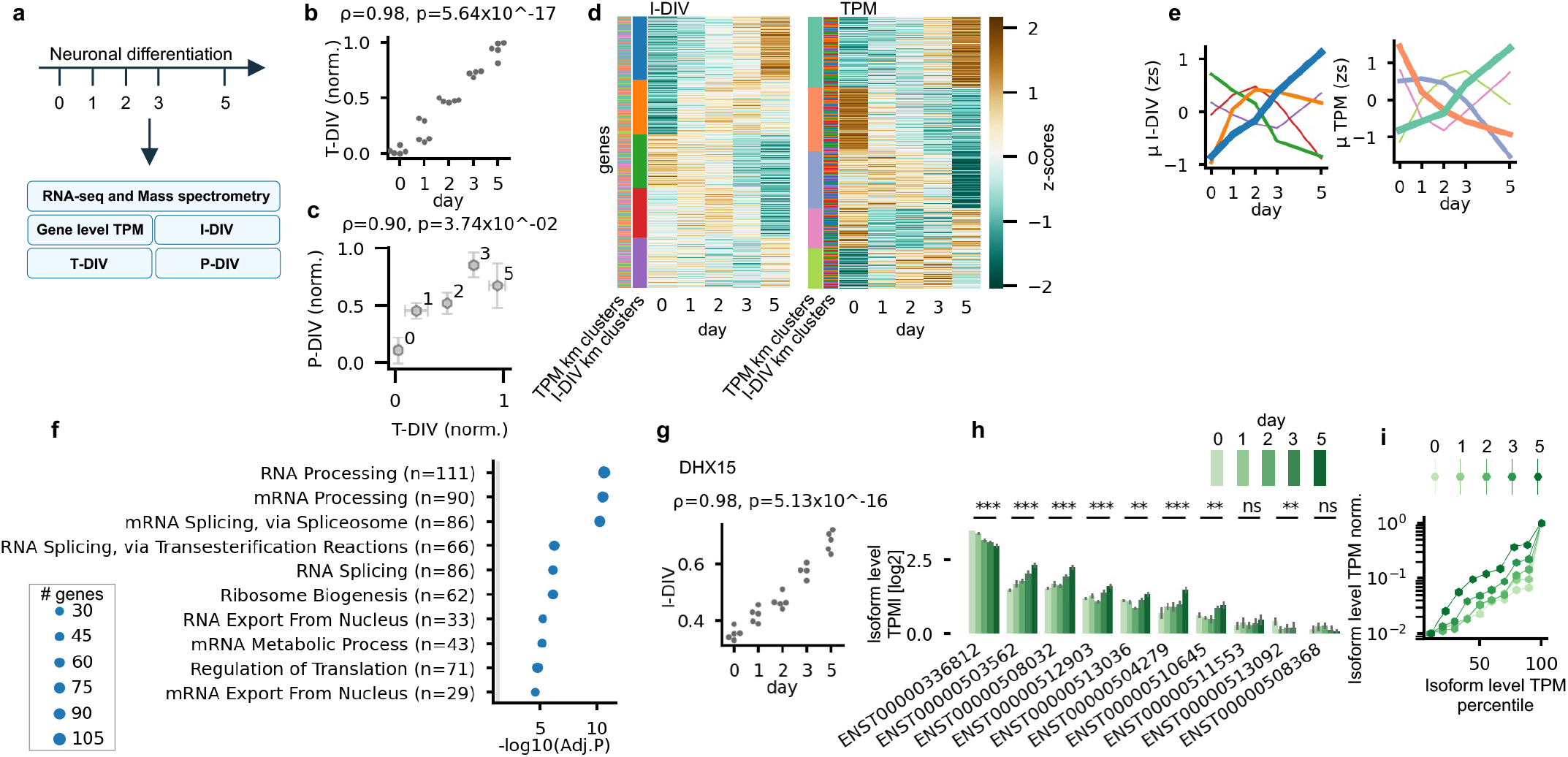
Isoform diversity changes during neuronal differentiation. (a) Schematic of the dataset used (Ulicevic et al., 2024), following a 5-day differentiation time course from iPSCs to neuronal cells, and the metrics calculated based on it. (b) T-DIV is shown by day of differentiation. Spearman’s coefficient and two-sided p-value are stated above. (c) T-DIV is plotted against protein diversity (P-DIV). Hexagonal datapoints represent mean of protein diversity and transcriptome diversity per day. Error bars show standard deviation: horizontal error bars for T-DIV; vertical error bars for P-DIV. Days are annotated near datapoints. Spearman correlation coefficient and p-value is stated above (T-DIV against P-DIV). (d) Heatmaps of mean I-DIV (left) and mean gene expression (right; TPM) per gene and per day of differentiation, z-scored. Each row corresponds to a gene and each column to a day of differentiation. Color bars indicate k-means clusters of gene expression (left) and of isoform diversity (right). Number of genes per I-DIV cluster, top to bottom (cluster A to E): 2,717, 2,371, 2,325, 2,162 and 2,152. Number of genes per gene expression cluster, top to bottom (clusters a-e): 3,047, 2,758, 2,460, 1,743 and 1,719. (e) K-means clusters show distinct trends. Genes were aggregated by k-means cluster of I-DIV or of gene expression, respectively. Colors correspond to the k-means cluster bars in panel d. Left: mean I-DIV of genes aggregated by k-means cluster of I-DIV. Right: mean gene expression of genes aggregated by k-means cluster. Y axis shown in z-scores. (f) Enrichment analysis using GSEApy enrichr of each I-DIV cluster (n = 2,717, 2,371, 2,325, 2,162 and 2,152 genes in clusters A to E) against a background of all genes in the analysis (n = 11,727), showing −log10(FDR) for the top 10 most highly enriched GO terms across all clusters. Colors correspond to the k-means cluster bars in panel d. The number of genes per term is indicated by size. (g) Swarmplot of *DHX15* I-DIV at different days of differentiation. Number of samples per day, in x-axis order: 5, 5, 5, 4 and 5; Spearman’s correlation coefficient and two-sided p-value is stated above. (h) Bar plot of mean expression of each *DHX15* isoform per day of differentiation. Error bars show the standard error; days are colored in shades of green. Significance of isoform changes measured in p-values from multiple linear regression (Methods): *** p < 0.001; ** p < 0.01; ns, not significant. (i) Line plot of the scaled mean expression of the ten *DHX15* isoforms, ordered by percentile of isoform expression across samples, for each day of differentiation (n=5, 5, 5, 4 and 5); days are colored in shades of green.

At the level of individual genes, isoform diversity and expression followed largely different trajectories (Fig. 5d), indicating a similar decoupling as observed in the GTEx dataset (Fig. 1). Clustering genes by their I-DIV profiles over time (k = 5; Supplementary Fig. 6c) identified one cluster of 2,717 genes whose I-DIV increased steadily throughout differentiation (blue line in Fig. 5de). Only this cluster was enriched for functional terms, predominantly RNA processing, mRNA splicing, RNA export and ribosome biogenesis (Fig. 5e). Clusters defined by expression did not correspond to the I-DIV clusters (Fig. 5c, TPM and I-DIV cluster are indicated for each gene using the color bars on the left) and, on the functional level, reflected the expected shift from proliferation and translation towards neuronal functions. Genes involved in cytoplasmic translation, DNA replication and ribosome biogenesis decreased, and genes involved in axon development, synapse organization and cilium assembly increased over the course of differentiation (Supplementary Fig. 6g–j), in line with the original analysis of day-to-day changes^48^. The splicing machinery itself therefore diversifies its isoform output as neurons form, consistent with RNA-binding proteins being key regulators of the widespread isoform changes during neurogenesis^48,50^. For example, I-DIV of the spliceosomal RNA helicase encoding gene *DHX15* increased from a median isoform diversity of 0.35 at day 0 to 0.69 at day 5 (Fig. 5g). Following the abundance of individual *DHX15* isoforms over time showed that the dominant isoform decreased while most minor isoforms increased (Fig. 5h), producing a progressively flatter isoform profile (Fig. 5i). The exon junction complex component encoding gene *RNPS1* showed a similar pattern (Supplementary Fig. 6d-f). Together with the GTEx analyses, these results show that isoform diversity is a dynamic and independent aspect of gene expression which captures regulatory changes across contexts, from tissue identity and ageing in a human cohort to cell-state transitions in a model of neuronal differentiation.

## Discussion

Gene expression changes at the mRNA isoform level are most often studied for individual splicing events or for differential isoform usage between conditions. By summarizing this distribution as a gene-level metric that is comparable across genes and samples, we observed that isoform diversity behaves as a gene property which can be decoupled from expression level. Across genes, I-DIV was almost uncorrelated with expression and only weakly associated with gene structure (Fig. 1b). Within genes, its coupling with expression was gene-specific and partly tissue-dependent (Fig. 1e). Isoform diversity separated tissues even based on genes with equivalent overall expression levels (Fig. 2). It was decoupled from transcriptome-wide diversity (Fig. 3), and it changed with age (Fig. 4) and during neuronal differentiation (Fig. 5), often without corresponding changes in expression.

I-DIV is related to other information-theoretic metrics of isoform usage^15,18^, in particular two recent approaches. SplicingFactory provides several gene-level indices, including normalized Shannon entropy and Laplace entropy with a fixed pseudo-count per isoform^21^. Perplexity expresses diversity as an effective number of isoforms and therefore increases with the number of isoforms a gene uses^22^. In contrast to previous approaches, I-DIV uses a pseudocount that scales with coverage, such that diversity values are comparable between samples with different depths of coverage and preventing lowly covered genes from being assigned spuriously high or low diversity values. Despite the differences between the metrics, perplexity applied to long-read sequencing data was likewise only weakly correlated with expression levels^22^, suggesting that this decoupling can be observed regardless of metric and sequencing technology.

Gene expression regulators and mRNA processing factors are intrinsically coupled to the generation of a pool of mRNA isoforms from the same gene, and a relationship between isoform diversity and the splicing machinery became evident in two independent context. In adult tissues, expression of genes involved in mRNA splicing, such as CDK12, correlated with mean isoform diversity across samples and tissues (Fig. 3j). During neuronal differentiation, RNA-processing and splicing genes were the only functional group whose own isoform diversity increased steadily (Fig. 5e). Ulicevic et al. observed that changes in the expression level of RNA-binding proteins were linked to isoform switches during neurogenesis. Our additional analyses suggest that not only the abundance, but also the functional diversification of splicing regulators are associated with the isoform diversity of their targets. Whether this reflects cause, consequence or a shared upstream program cannot be resolved from correlative analyses alone and will require perturbation experiments.

A change in isoform diversity describes how a gene’s expression is redistributed across its isoforms. An increase in I-DIV means that expression is spread more evenly, often because alternative isoforms become more abundant at the expense of the dominant one. This can broaden the repertoire of gene products and may point to functional diversification. A decrease in I-DIV implies that the gene’s expression becomes concentrated on fewer isoforms, which may reflect commitment to a specific product. In the case of bulk RNA-seq samples, changes in I-DIV can arise in different ways: through regulatory changes in splicing or transcript start and end site selection, through changes in cellular composition, since mixtures of cell types with different dominant isoforms appear more diverse, or through technical noise in isoform quantification. Interpreting a change therefore requires inspecting the underlying isoforms, ultimately on a single-cell level. As a gene-level summary, I-DIV complements isoform-by-isoform analyses of differential transcript usage^51^. Importantly, I-DIV cannot be used to identify switches between isoforms that leave the overall distribution unchanged. However, it requires one test per gene rather than per isoform, which reduces the limitations of multiple testing. In addition, it can capture coordinated shifts across many minor isoforms that may each be too small to reach significance on their own.

Age-related changes in the transcriptome have often been interpreted as a loss of regulatory precision, through increasing heterogeneity between individuals or cells^36,38,52^. In the GTEx dataset, age-related changes in gene expression have been shown to differ between tissues^31^ and between the sexes^33^, and are accompanied by tissue-specific changes in alternative splicing^34,35^. Our results add an additional dimension of gene expression diversification. Age was associated with isoform diversity in a gene- and tissue-specific manner, most strongly in the brain, where mean I-DIV decreased with age in most regions while transcriptome diversity showed limited changes. This suggests a general trend where isoform usage in the ageing brain shifts towards fewer dominant isoforms. In contrast, I-DIV in a set of genes increased with age, including *PPARD*, in which isoforms lacking the DNA-binding domain or carrying an alternative N-terminal disordered region increased with age at the expense of the dominant isoform. Whether such shifts alter protein function remains to be tested, but they illustrate how changes in isoform diversity can point to functional changes that an analysis of expression levels alone may miss.

Several conceptual and technical limitations apply. Firstly, isoform diversity is measured at the RNA level. Not every mRNA isoform is translated or exhibits different functional properties^13^, so I-DIV is a proxy for, not a measure of, functional diversification at the protein level. However, in the absence of extensive protein isoform-resolved proteomics data, I-DIV can provide starting points for investigating specific hypotheses regarding the functional consequence of changes in the isoform composition, such as in the case of *PPARD*. As mentioned before, functionally relevant isoform switches would not be detectable in an I-DIV based analysis if they did not change the overall distribution of isoforms, emphasizing that gene-level diversity analyses should be seen as complementary to existing analytical approaches. Lastly, while we and others^22^ observed overall decoupling of gene expression level and isoform diversity, they were correlated within many individual genes, positively for some and negatively for others (Fig. 1e). Our dynamic pseudocount reduces, but cannot eliminate, the dependence of I-DIV on sequencing depth, as isoform proportions of poorly covered genes are estimated from few reads. In cases where a gene’s expression, and consequently its coverage, differs strongly between conditions, differences in I-DIV should be interpreted together with the underlying isoform-level expression to determine whether it reflects a genuine redistribution of expression between isoforms rather than an artifact related to minimal coverage.

Taken together, our results show that the diversification of a gene’s expression on the mRNA level is a distinct and complementary dimension of gene expression. It can vary with tissue identity, age and cell state, often independently of expression level and overall transcriptome diversity. By providing a robust gene-level metric for isoform diversity, I-DIV complements differential gene or isoform expression analyses. It constitutes a quantitative trait that can be linked to genetic variation or disease and has the potential to identify changes in a gene’s functional output.

## Methods

### Data sources

We used GTEx v8 RNA-sequencing data (release 2017-06-05): gene-level TPM (RNA-SeǪC v1.1.9) and transcript-level TPM (RSEM v1.3.0). Sample attributes and subject phenotypes (sex, age bracket and death classification on the four-point Hardy scale) were taken from the GTEx v8 annotation files. Gene and transcript annotation was taken from GENCODE v26 (GFF3), which matches the annotation used for GTEx v8. Tissues were analysed at two levels: SMTS (broad tissue type) and SMTSD (tissue subtype, e.g. individual brain regions). Each SMTSD was also assigned to an organ system (e.g. nervous, digestive, cardiovascular).

### Sample filtering

Sample and donor metadata were merged by donor identifier. We retained only samples from donors with a Hardy-scale death classification of 1 (violent and fast death) or 2 (fast death of natural causes), to minimise effects of prolonged illness or terminal state on expression.

Missing RNA integrity numbers (RIN) and missing ischaemic times (SMTSISCH) were imputed with the mean of the samples from the same SMTSD, and negative ischaemic times were set to zero. Only samples with RIN ≥ 6 were retained. Only SMTSDs with at least 15 remaining samples were kept. Finally, samples were retained only if present in all four GTEx expression matrices (gene counts, gene TPM, transcript counts, transcript TPM).

The final dataset comprised 4,814 samples from 267 donors, 25 SMTS tissues, 47 SMTSD tissues, including 13 brain SMTSD regions.

### Gene filtering and gene annotation

Genes were restricted to protein-coding genes, excluding chromosomes X, Y and M, to avoid sex-linked and mitochondrial effects. 18380 genes were retained after this filtering step. Gene length was calculated as end minus start coordinate, and the number of exons per gene was counted from the non-redundant exons of the GENCODE annotation. The number of potential isoforms of each gene was defined as the number of its transcripts in the GTEx transcript matrix.

Ribosomal protein (RP) genes were defined as all genes annotated to any Gene Ontology Cellular Component 2025 term (Enrichr library, downloaded with gseapy) whose name contains “ribosom”. Each gene was then labelled as RP.

A second filter was applied after the isoform diversity calculation (see below): genes were kept only if more than 1% of samples had an isoform diversity above zero, implying that it had non-zero expression of at least two isoforms in more than 1% of samples. This filter was applied to every matrix used downstream (gene-level TPM, isoform diversity, total expression, number of expressed isoforms, gene metadata), resulting in a final set of 15,834 genes, including 144 ribosomal protein genes .

### Isoform diversity (I-DIV)

To quantify how evenly a gene’s expression is distributed across its annotated isoforms, we calculated an isoform diversity index (I-DIV) for each gene in each sample from transcript-level TPM values. I-DIV is an adaptation of Shannon’s evenness index, which divides the Shannon entropy of a distribution by its maximum possible value.

For a gene with *n* annotated isoforms, let *x*_1_, … , *x_n_* be the expression values (TPM) of its isoforms in a given sample, and let 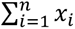 be the gene’s total expression. If at most one isoform had non-zero expression, I-DIV was set to 0 (this includes all genes with a single annotated isoform). Otherwise, I-DIV was calculated in three steps.

Firstly, a dynamic, gene- and sample-specific (dynamic) pseudo count *d* was calculated as follows:

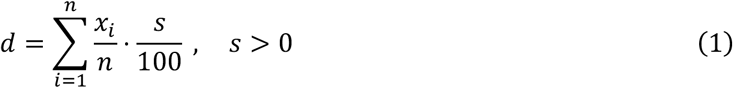

*d* is calculated as the percentage *s* of the mean expression level of an (annotated) isoform of a gene (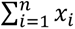 divided by *n*). *s* can be varied and was usually set to 1 (percent) in this study, unless stated otherwise. In other words, typically 1% of a gene’s total expression in a sample was divided equally among its *n* annotated isoforms. This approach can be used both on counts data and on expression estimates (TPM, CPM, etc.).

Secondly, this dynamic pseudocount was added to the expression value of every annotated isoform.

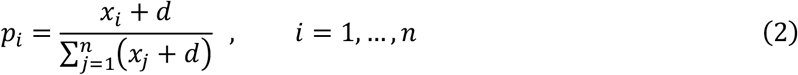

*p_i_* denotes the proportion of isoform *i* expression with an additional value of a dynamic pseudocount (*d*). As a result, isoforms that are annotated but not expressed receive a small, non-zero proportion and are included in the entropy calculation (see below, Eq. 3).

Thirdly, Shannon’s entropy was calculated based on the proportions including the dynamic pseudocount:

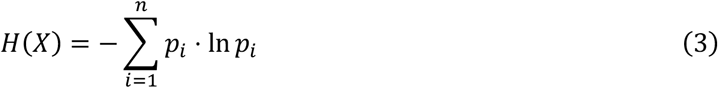

Lastly, the entropy *H* of the isoform-level expression distribution of a gene as a whole (*X*) was divided by the natural logarithm of the number of annotated isoforms (n) to calculate a metric for isoform diversity (I-DIV):

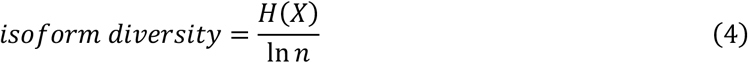

I-DIV ranges from 0 to 1, where 0 indicates the presence of only one annotated (and expressed) isoform and 1 implies equal (non-zero) expression of all (>1) annotated isoforms. Because *d* is proportional to the expression level of the gene, I-DIV depends only on the relative expression of a gene’s isoforms, not on the gene’s overall expression level.

### Transcriptome diversity (T-DIV)

Transcriptome diversity was calculated per sample by applying the same diversity function (1% dynamic pseudocount) to the vector of gene-level TPM values (for the indicated gene groups after filtering), so that it captures how evenly expression is distributed across the transcriptome. Throughout the paper, T-DIV refers to all genes except the ribosomal protein genes unless stated otherwise.

### Cellular composition diversity (cell-DIV)

To compare transcriptome-wide diversity with the diversity of cell types in a sample, we used the cellular composition estimates that Donovan et al.^29^ calculated for GTEx samples. They deconvoluted bulk GTEx RNA-seq samples using CIBERSORT, a quantitative tool to estimate the abundances of cell types in mixed cell populations. Supplementary Data 4-8, 10-14 and 17 were used here, including aorta, heart (atrial appendage and left ventricle), the 13 brain regions, colon (sigmoid and transverse), adipose tissue (subcutaneous and visceral), liver, breast, skeletal muscle, pancreas and skin (sun exposed and not sun exposed). Cellular diversity (cell-DIV) of each sample was calculated as the Shannon entropy of its cell-type proportions.

### Global comparison of isoform diversity with gene features

For every gene we calculated, across all 4,814 samples, the mean log2(TPM+1), the mean I-DIV and the mean expressed isoform count per gene. Genes with more than two exons, a mean expressed isoform count per gene above 2 and a mean I-DIV above 0 were kept (11,981 genes). Across these genes, mean I-DIV and mean log2(TPM+1) were each correlated (SciPy pearsonr) with the number of exons, gene length (data was obtained fro GENCODE gff file version 26), the number of potential isoforms and mean expressed isoform count, and mean I-DIV was also correlated with mean log2(TPM+1).

### Tissue-specific comparison of isoform diversity to gene level expression

For every gene we correlated (SciPy pearsonr) I-DIV against gene exprression log2(TPM+1), and I-DIV against number of expressed isoforms, separately within each non-sex-specific SMTS (19 tissues; Uterus, breast, testis, ovary, vagina and prostate are excluded). For a given gene, a gene with at least 10 samples whre I-DIV > 0 in a tissue to be tested; otherwise its coefficient was set to 0 and its p-value to 1. The per-tissue Pearson coefficients of I-DIV against gene expression formed a matrix of 15,834 genes × tissues. Genes were clustered with k-means (scikit-learn KMeans) on the coefficient matrix.

### Gene-set enrichment analysis

Over-representation analysis was run with Enrichr through gseapy (enrichr function) on the GO Biological Process 2025 library. In every analysis a custom background was supplied which contained all genes that were used in the analysis, as specified in each section. Enrichr p-values are from Fisher’s exact test and were adjusted with the Benjamini–Hochberg method within each library and query; terms with adjusted p < 0.05 were considered enriched.

### Pairwise differential expression/diversity and equivalence testing between tissues

Fitting limma models to pairwise groups of tissue samples

To identify genes whose expression or diversity differs between two tissues, and genes for which it does not, we compared every pair of tissues, at the SMTS level (25 tissues, 300 pairs) and at the SMTSD levels within Brain samples (13 brain regions, 78 pairs). Differences were tested with limma (lmFit and eBayes) using a design with tissue as the only term. P-values were adjusted for multiple testing across genes with the Benjamini– Hochberg method. The estimated effect (Δ; Change) is the difference in mean (log2(TPM+1) or I-DIV) between the two tissues.

Identifying non-differentially expressed genes

Equivalence was tested with a two one-sided tests (TOST) procedure on the same fitted model, with an equivalence margin of δ = 0.5. For each gene we tested that the difference is greater than −δ and that it is smaller than +δ, using the standard error from the linear model and the moderated degrees of freedom from eBayes. The larger of the two one-sided p-values was taken as the equivalence p-value and adjusted across genes with the Benjamini– Hochberg method. A gene was called equivalent when its adjusted equivalence p-value was below 0.05. Every gene in every pair was then assigned to one of three classes for gene expression: differentially changed (adjusted p < 0.05 in the difference test), equivalent (not differentially changed, adjusted equivalence p < 0.05), or inconclusive (neither). A gene that differed significantly was therefore never called equivalent. The analysis was implemented in R 4.3.3 with limma 3.58.1 and data.table with one job per tissue pair.

Principal component analysis and silhouette scores

To measure how well gene expression and isoform diversity each separate two tissues, we ran, for every tissue pair, a principal component analysis on the samples of the two tissues. Each gene was standardised across these samples (scikit-learn StandardScaler), the first two principal components were computed (scikit-learn PCA), and the separation of the two tissues was measured with the silhouette score of the tissue labels in the PC1–PC2 space (scikit-learn silhouette_score). This was done separately on log2(TPM+1) and on I-DIV, and with two gene sets: all 15,834 genes, and only the expression-equivalent genes between the two tissues (with mean log2(TPM+1) above 0, limma AveExpr). Silhouette scores of expression and of I-DIV were compared across pairs with the Wilcoxon signed-rank test (SciPy wilcoxon), for all genes and for expression-equivalent genes, separately for all the pairs described above.

Differential expression and diversity analysis

We used the limma results above and kept only genes with a median I-DIV above 0 across the samples of the pair. Genes were classified as having significantly changed expression in case of adjusted p < 0.001 and Δ mean log2(TPM+1) ≥ 1. Genes were classified as having significantly changed diversity in case of adjusted p < 0.05 and Δ mean I-DIV ≥ 0.05. For each pair and direction, enrichment was tested as described above, with all genes used for the analysis of each pair as background. Terms with adjusted p < 0.05 and at least 5 genes were reported for expression and I-DIV.

### Correlation analysis of individual gene expression levels with I-DIV and T-DIV

To find genes whose expression is associated with transcriptome diversity or mean isoform diversity, we correlated the expression of every gene with T-DIV and with µ I-DIV across samples, for each non-sex specific SMTS tissue (19 tissues) separately. Gene expression values were transformed to log2(TPM+1) and then z-scored within each sample across genes (SciPy zscore), so that each value expresses how a gene is expressed relative to the other genes in the same sample.

For each tissue, we calculated the Pearson correlation for every gene with TPM>0 in at least 10 samples between its z-scored expression and T-DIV, and separately µ I-DIV. For genes with TPM>0 in less than 10 samples in a tissue its coefficient was set to 0 and its p-value to 1. For T-DIV, 1% of samples with the lowest number of expressed genes were excluded. This filter was not applied for µ I-DIV.

The analysis used the 19 non-sex-specific SMTS tissues. Within each tissue, p-values were adjusted across genes with the Benjamini–Hochberg method (statsmodels multipletests, untested genes excluded). A gene was assigned to a positively (or negatively) correlated group of a metric if its correlation was significant (FDR < 0.05) with the same direction of correlation in at least 10 of the 19 tissues, yielding four groups: µ I-DIV+, µ I-DIV−, T-DIV+ and T-DIV−. Each group was tested for enrichment of GO Biological Process 2025 terms, with all 15,834 genes as background. Terms with fewer than 10 genes from the query were not considered, and terms with adjusted p < 0.05 were labeled enriched. Enriched terms were classified as enriched for T-DIV only, for µ I-DIV only, or for both, separately for positive and negative correlations.

### Identification of age- and sex-associated changes in I-DIV

#### Age increments mapping

To measures the change of age per tissue with more balanced age groups, we grouped the samples according to three age brackets: 20-49, 50-59, 60-79. In linear mixed model and multiple linear regression tests, the three age groups were represented by 1,2,3, respectively.

#### Association of age and sample-level metrics

The association of age with each metric was measured with Spearman’s rank correlation coefficient (SciPy spearmanr). For each of the 25 SMTS and 13 brain SMTSD tissues, the coefficient (ρ) and its two-sided p-value were calculated between the metric and the three age groups across all samples per tissue. This was done separately for T-DIV and for µ I-DIV.

#### Ǫuantile normalization

For the gene-level analyses, gene expression and isoform diversity were quantile normalized separately within each tissue, at the SMTS level (25 tissues) or at the SMTSD level (47 subtissues), and the tissues were then combined into one matrix per measure. Within a tissue, the values of each sample were ranked across genes, and tied values received their average rank. The reference distribution was the mean, over the samples of the tissue, of the sorted values at each rank. Each value was replaced by the reference value at its rank, interpolating linearly for tied ranks. After this step every sample of a tissue has the same distribution of values. Gene-level TPM values were transformed to log2(TPM+1) and quantile normalized within each tissue. For I-DIV, transcript-level TPM of all transcripts were first quantile normalized within each tissue. I-DIV with 1% dynamic pseudocount was then calculated for every gene and sample from the quantile normalized transcript values.

#### Linear mixed effects models

To estimate an age- and sex-associated variation shared between tissues, we used linear mixed models with the indicated fixed effects and donor as a random effect (because each donor contributes samples to several tissues) and fitted a model for each gene, following Melé et al., 2015. In contrast to Melé et al., 2015, we used quantile normalized values as described above. Three variations of the linear mixed models were run using nlme::lme (version 3.1.171) in R:

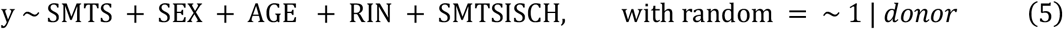

All SMTS tissues (25 tissues) together, with sex, RIN and ischemic time as covariates (formula 5; Fig. 4e left, Supplementary Fig. 5c bottom, 5d).

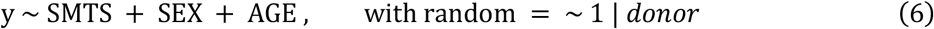

The same model as formula 5, without RIN and ischemic time to compare genes with significant age coefficients with the work from Melé et al. 2015^32^ (formula 6; Supplementary Fig. 5c top);

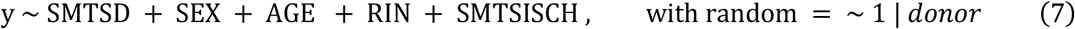

Applied on 13 brain SMTSD regions together, with sex, RIN and ischemic time as covariates (formula 7; Fig. 4e right, Supplementary Fig. 5e).

In all three formulas (5, 6, 7), y denotes the response variable, i.e. gene expression (log2(TPM+1)) or I-DIV. Models were fitted by restricted maximum likelihood with the lme function. For each term of the equation, the coefficient, standard error, degrees of freedom, t-value and p-value of the fixed effects were computed, and p-values were adjusted with the Benjamini–Hochberg method across genes, separately for each term. Genes were called associated with a certain term when the FDR of the term was below 0.05.

#### Comparison with a published age analysis

The age coefficients of the all-tissue mixed model for gene expression were compared with those of Table S12 of Melé et al. 2015^32^ (genes matched by Ensembl ID without version number). Pearson’s correlation was calculated over the genes that were significant in this study (FDR < 0.05) and also over all matched genes. The comparison was repeated for the mixed model without RIN and ischemic time, which is closer to the published model.

#### Multiple linear regression of age, sex and technical covariates per tissue on the gene level

Following García-Pérez et al., 2023^35^, per-tissue effects were estimated with limma. The effect of age on each gene was estimated with a multiple linear regression fitted separately within each tissue, using limma 3.58.1. The response was either gene expression or the I-DIV of the gene. Each analysed on its own. Tissues were analysed at the SMTS and at the SMTSD level (25 SMTS tissues, 47 SMTSD respectively).

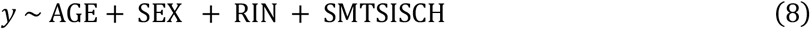

For each gene, the response was regressed on age, sex, RIN and ischemic time. In SMTS-level runs, the tissue subtype (SMTSD) was added when the tissue has several subtypes. Age was coded as an ordinal variable with three groups, 20–49 = 1, 50–59 = 2 and 60–79 = 3, so the age coefficient is the mean change per step to the next group. Sex was a two-level factor, and RIN and ischemic time were numeric. Covariates without variation in a tissue, for example a single sex, were dropped, and samples with a missing covariate were excluded.

Models were fitted with lmFit and eBayes with the trend option, so the variance of each gene is moderated towards a prior that depends on its mean level. P-values were adjusted with the Benjamini–Hochberg method across the tested genes of a tissue, separately for each term. Genes were called age-associated when the FDR of the age term was below 0.05.

#### Gene-level Spearman correlations with age per tissue

Age was compared with isoform diversity and gene expression level using. Spearman’s coefficients and P-values were obtained. P-values were djusted by the Benjamini–Hochberg procedure within each tissue and separately for gene expression and I-DIV. A gene was called age-associated when corrected FDR was below 0.05.

#### Multiple linear regression of age, sex and technical covariates per tissue on the isoform level

To test how the expression of individual isoforms changes with age, the same regression was fitted (formula 8) per isoform for selected genes (PPARD, FAM161B and KNDC1). For each gene, all of its isoforms were analysed in each brain SMTSD, one tissue at a time.The data were transcript-level TPM quantile normalized within each brain SMTSD (see above), The model, the age coding and the handling of covariates are the same as in the gene-level regression. The asterisks in the figure mark isoforms by the unadjusted p-value of the age term, because statistical testing was done only on isoforms of the same gene and individual isoform-level tests are not indipendent.

### Analysis of isoform diversity changes during neuronal differentiation

#### Dataset

To follow isoform diversity during neuronal differentiation, we used the published RNA-seq and proteomic data of Ulicevic et al. (Ulicevic J, Shao Z, Jasnovidova O, et al., Mayer A. Uncovering the dynamics and consequences of RNA isoform changes during neuronal differentiation. Mol Syst Biol. 2024;20(7):767–798; doi:10.1038/s44320-024-00039-4). We used the RSEM quantification provided with the study, based on GENCODE v28: gene-level and transcript-level TPM for every sample, with samples named by day of differentiation and replicate (RSEM v.1.2.25; human reference genome GRCh38 version p12).

#### I-DIV calculation

I-DIV (dynamic pseudocount of 1%, as for GTEx) and number of isoforms were calculated per gene and sample from the transcript-level TPM. Genes were restricted to protein-coding genes outside chromosomes X, Y and M (GENCODE v28) which results in 18,984 genes. Using a cutoff of I-DIV > 0 in at least 80% of samples resulted in a final number of genes used of 11,727. 84 RP genes were flagged, using gseapy, Biomart and GENECODE version 26.

#### T-DIV calculation

Per sample we calculated transcriptome diversity (T-DIV) as described above, excluding ribosomal protein genes. To allow for a larger dynamic range in the display of the resulting varlues, we applied min-max scaling across the T-DIV values.

#### Protein diversity (P-DIV) calculation

Protein diversity (P-DIV) was calculated from the MaxǪuant proteinGroups table of the study’s PRIDE project (PXD046084, isoform_txt.zip). Protein groups whose gene names matched the genes used for T-DIV were kept, and P-DIV of each sample was calculated with the same diversity function (1% dynamic pseudocount) over the “Unique peptides” counts of these protein groups, then min–max normalised across samples.

#### Clustering analyses

For identifying common trends of changes in gene expression or isoform diversity across neurodifferentiation, log2(TPM+1) and I-DIV were z-scored per gene across samples, averaged per day, and clustered separately with k-means. One gene (ENSG00000145386) was dropped because its I-DIV was the same in all samples. Each cluster was tested for enrichment against all clustered genes.

#### Individual gene analyses

For the example genes DHX15 and RNPS1, I-DIV was correlated with day (Spearman), and the expression of each isoform was compared between day 0 and day 5 using differential expression with limma. For the isoforms of the tested gene, TPM values were transformed as log2(TPM + 1). A linear model with day (day 5 versus day 0) as the only term was fitted to each isoform with lmFit, treating replicates as independent samples with no pairing. Variances were moderated with the empirical Bayes procedure eBayes, without a mean-variance trend. For each isoform, the log2 fold change (day 5 relative to day 0), the moderated t-statistic and the p-value were obtained with topTable. P-value thresholds: * ≤ 0.05, ** ≤ 0.01 and *** ≤ 0.001 .

### Implementation

Analyses were implemented in Python 3.12.2 with pandas 2.2.2, NumPy 1.26.4, SciPy 1.13.0, statsmodels 0.14.1, scikit-learn 1.3.0, gseapy 1.1.4, mygene 3.2.2, seaborn 0.13.2 and matplotlib 3.8.4, isoplex 0.1.0. Large matrices were read in chunks of selected rows, and per-gene calculations were parallelised with SLURM array jobs on a high-performance computing cluster. The GENCODE v26 and v28 annotations were downloaded from the GENCODE FTP site and parsed with gffpandas 1.2.2. R packages nlme 3.1.171 and limma 3.58.1 were run in R 4.3.3.

### Computational infrastructure used

The pipeline was executed on the HIVE high-performance computing cluster at the University of Haifa with: Red Hat Enterprise Linux 9 operating system distribution, SLURM job scheduling for parallel processing (v23.02.4)

## Data availability

All analyzed data are publicly available. Gene- and transcript-level expression data (TPM) and sample and subject annotations from the Genotype-Tissue Expression (GTEx) project release v8 are available from the GTEx Portal (https://www.gtexportal.org/home/downloads/adult-gtex/). RNA-seq and mass spectrometry data from the human iPSC-derived neuronal differentiation time course (Ulicevic et al.^48^) were obtained from the Gene Expression Omnibus (accession GSE245325) and the PRIDE database (accession PXD046084). Cell-type proportions for GTEx samples are available as Supplementary Data accompanying Donovan et al.^29^

## Code availability

Computational methods and custom scripts are available in a GitHub repository: https://github.com/shakedshanas/I-DIV_dim

## Author contributions

S.S., E.L. and S.J. performed analyses, S.S., E.L. and M.M. conceived the project, S.S. and M.M. wrote the manuscript.

