## Supplementary Figures for "Isoform diversity as a complementary dimension of gene expression"

Supplementary Figure 1: Simulation of the effect of coverage on isoform diversity estimates.

Supplementary Figure 2: Isoform diversity is largely decoupled from other gene features.

Supplementary Figure 3: Enrichment analysis of genes differing in expression or isoform diversity between cerebellum and cortex.

Supplementary Figure 4: Identification of genes whose expression is significantly and robustly associated to T-DIV or μ I-DIV.

Supplementary Figure 5: Changes in isoform diversity and gene expression with age.

Supplementary Figure 6: Additional analyses of neuronal differentiation.

Supplementary Data

Supplementary Data 1: k-means cluster of genes by I-DIV to TPM correlation profiles.

Supplementary Data 2: Pairwise separation ability of samples in brain subregions based on all genes and non-differentially expressed genes.

Supplementary Data 3: GO term enrichment analysis for genes whose expression is significantly and robustly associated to T-DIV or μ I-DIV.

Supplementary Data 4: Coefficients and p-values for linear mixed models of I-DIV and expression level for each gene in brain.

Supplementary Data 5: Associations of I-DIV and TPM with age, using limma.

Supplementary Data 6: Associations of I-DIV and TPM with age, using Spearman.

### Supplementary Figures

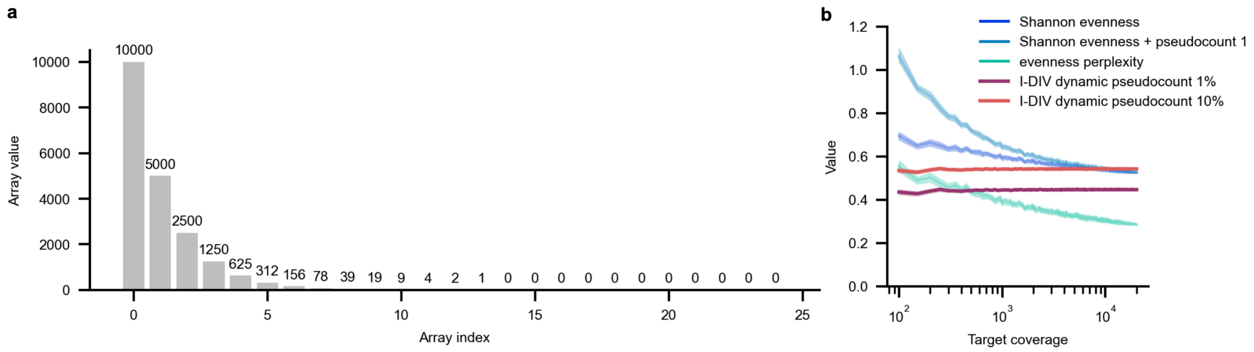


Supplementary Figure 1: Simulation of the effect of coverage on isoform diversity estimates.

(a) Barplot describing the underlying distribution of isoform abundances of a gene from which an RNA-sequencing experiment samples. This distribution represents an exponential series of 25 values, values decrease by 10,000 / $2^k$ for k = 0-24 (14 non-zero values from 10,000 down to 1, and 11 zeros; in total 19,995 counts). (b) Different diversity metric scores plotted against the (simulated) coverage of the target gene from panel a. From the constant underlying distribution (panel a) we iteratively subsampled without replacement to target coverage (x-axis, 30 repetitions per target coverage) and calculated the resulting isoform diversity based on different metrics. The diversity metrics used here were Shannon evenness (dark blue), Shannon evenness with a constant pseudo count of 1, the evenness metric based on perplexity values as described in Schertzer et al., 2026 (turquoise), isoform diversity with dynamic pseudocounts (I-DIV, this study) based on 1% (purple) and 10% (orange) of the total gene coverage distributed among all potential isoforms. The target coverage (sum of the array) increased from 100 to 19,950 in steps of 50, in addition to calculation of diversity metrics on the full depth of 19,995, resulting in 399 levels of coverage and 11,970 iterations in total.

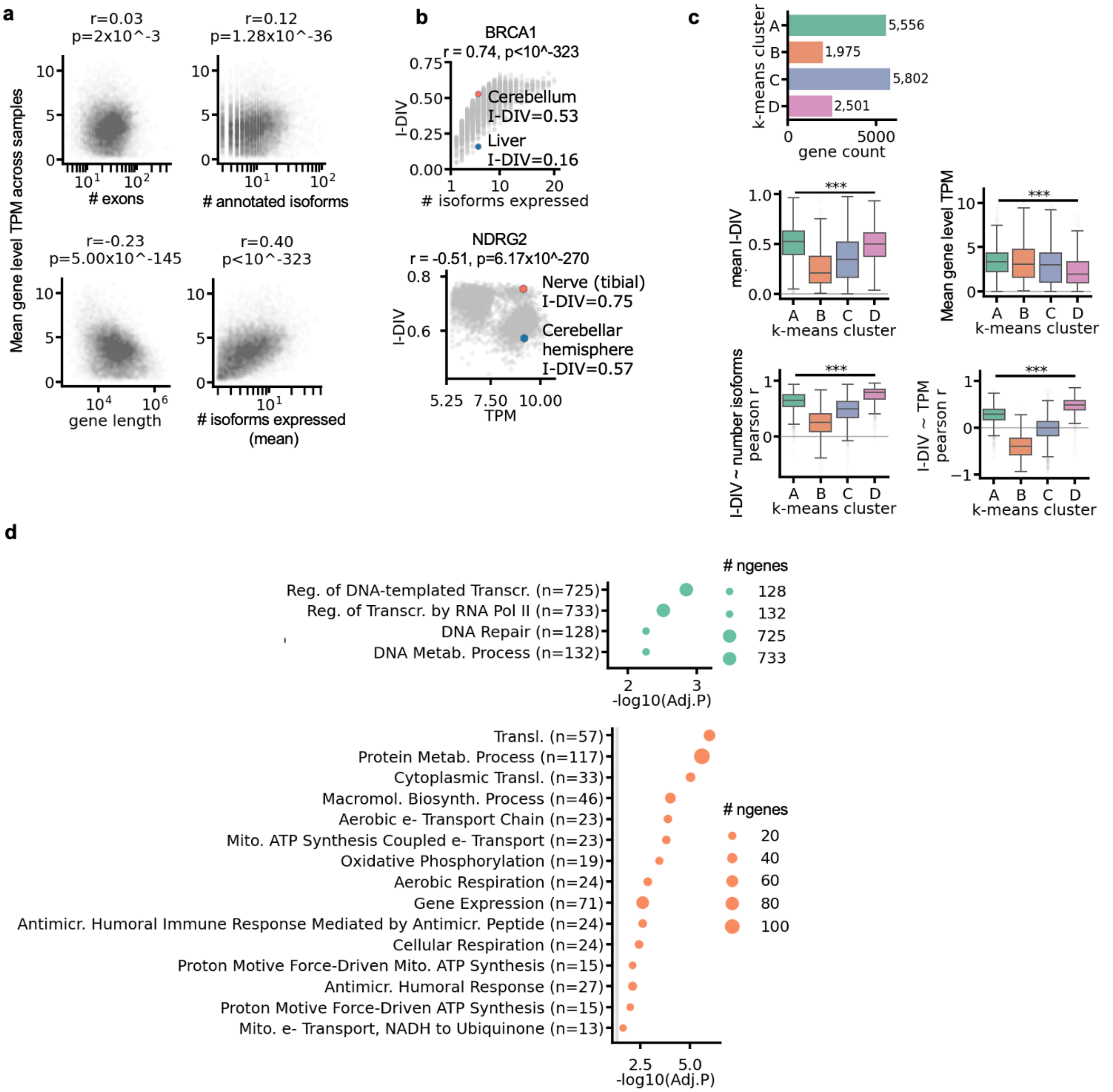


Supplementary Figure 2: Isoform diversity is largely decoupled from other gene features.

(a) Overview of gene level log2(TPM+1) (TPM) in comparison with gene features. Every datapoint represents a different gene (n=11,981), mean measurements are across 4814 GTEx samples. Top panels, left to right: mean TPM plotted against number of exons (# exons), number of annotated isoforms (# annotated isoforms) . Bottom panels, left to right: TPM plotted against gene length, and mean number of expressed isoforms (# isoforms expressed). Pearson's Correlation coefficients and p-values are stated in each panel. (b) *BRCA1* and *NDRG2* genes. Top: *BRCA1* I-DIV plotted against number of isoforms expressed (n = 4,813; Pearson r = 0.3, p =$1.21x10^{-101}$). Highlighted hexagons mark a cerebellum sample colored in salmon (I-DIV=0.53; # isoforms expressed= 6) and a liver sample in blue (I-DIV=0.16; # isoforms expressed=6), each expressing six isoforms. Bottom: *NDRG2* I-DIV plotted against TPM. Highlighted hexagons mark a nerve sample in salmon color (I-DIV=0.75;

log2(TPM+1)=9.15) and a cerebellar hemisphere sample in blue (I-DIV=0.57; log2(TPM+1)=9.19) Pearson's correlation coefficient and two-sided p-value are stated above. (c) Evaluation of the k-means gene clusters. The bar plot shows the number of genes per cluster. Box plots show the median (central line) and interquartile range (box, $25^{th}$–$75^{th}$ percentiles), with whiskers extending to the most extreme data point within 1.5x the interquartile range; outliers are shown as light grey points. One-way ANOVA for differences between clusters is reported for each panel. Box plots are shown in the following order: Mean I-DIV (One-way ANOVA $p < 10^{-323}$); mean gene expression (One-way ANOVA $p = 3.20 \times 10^{-121}$); mean Pearson's correlation coefficients between I-DIV and gene expression across tissues (One-way ANOVA $p < 10^{-323}$); mean Pearson's correlation coefficients between I-DIV and the number of expressed isoforms (One-way ANOVA $p < 10^{-323}$); and mean correlation between I-DIV and gene expression (log2(TPM+1)) (One-way ANOVA $p < 10^{-323}$). Colors correspond to the clusters in Fig. 1 panel e. (d) Enrichment analysis using GSEApy enrichr of clusters A (top; n = 5,556) and B (bottom; 1,975) against a background of all genes in the analysis (n = 15,834). Showing -log10(FDR) for the top 10 most highly enriched GO terms across all clusters. Colors correspond to the k-means cluster bar in Fig. 1 panel e. The number of genes per term is indicated by size.

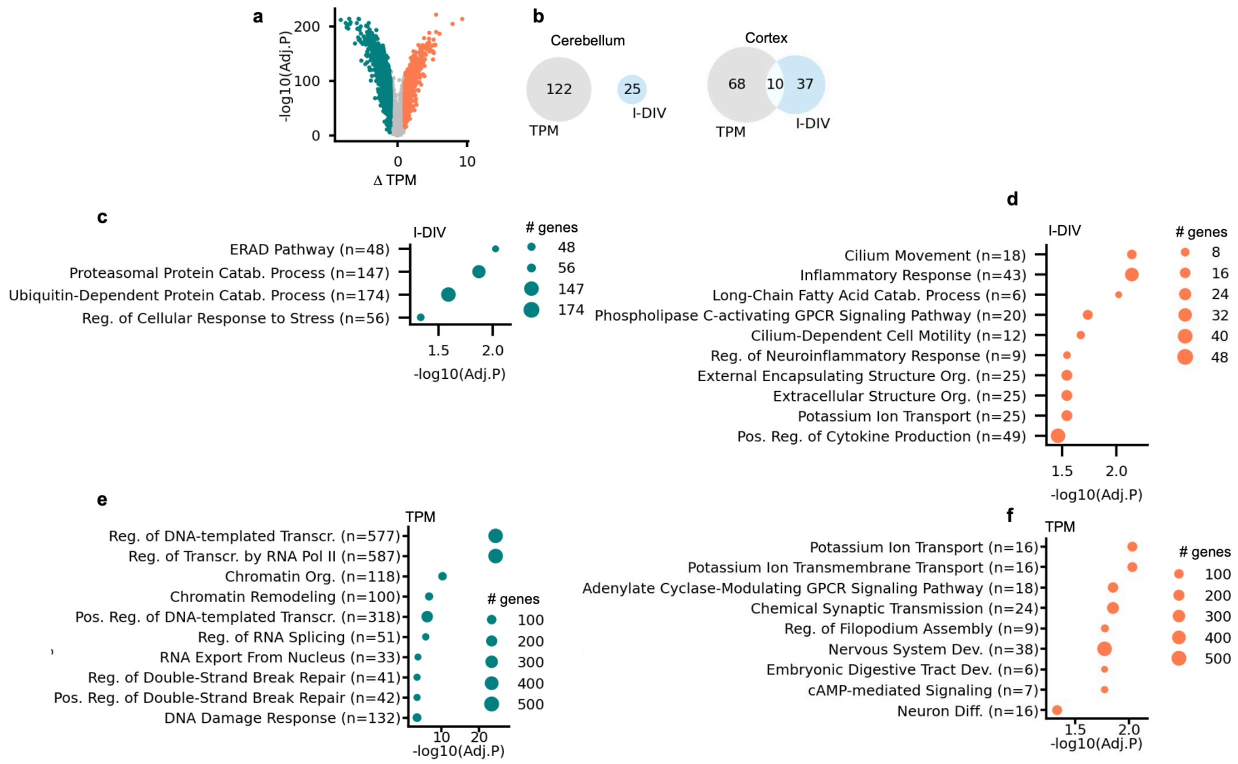


Supplementary Figure 3: Enrichment analysis of genes differing in expression or isoform diversity between cerebellum and cortex.

(a) Volcano plot of change in gene expression (cortex-cerebellum) against the p-value corrected for multiple testing (tested using limma, FDR, Benjamini–Hochberg). Genes with median I-DIV above 0 across cortex and cerebellum samples were used (n = 13,328). Cut-offs for significance were adjusted $p < 0.001$ and an absolute log fold change ≥ 1. Teal colored: genes with higher expression in cerebellum (n = 3,411); Coral colored: genes with higher expression in cortex (n = 717). (b) Venn diagrams showing Number of enriched terms of genes changed in cortex and cerebellum. Venn diagrams showing the number of enriched terms (GSEApy Enrichr; FDR < 0.05 and at least 5 genes per term) per metric. Query gene lists were genes with positive change of gene expression (grey) or change of I-DIV (light blue). Left: number of enriched terms by genes with higher values in cerebellum. Right: number of enriched terms by genes with higher values in cortex. Size of circles indicate number of terms. (c-f) Enrichment analysis using GSEApy enrichr of genes with higher I-DIV in cortex (n=1,853, 5,115, 717, 3,411 respectively) a background of genes with median I-DIV above 0 across cortex and cerebellum samples (n = 13,328). The number of genes per term is stated. Enrichment threshold is by corrected p-values for multiple testing (FDR, Benjamini–Hochberg method) below 0.05. The number of genes per term is indicated by size.

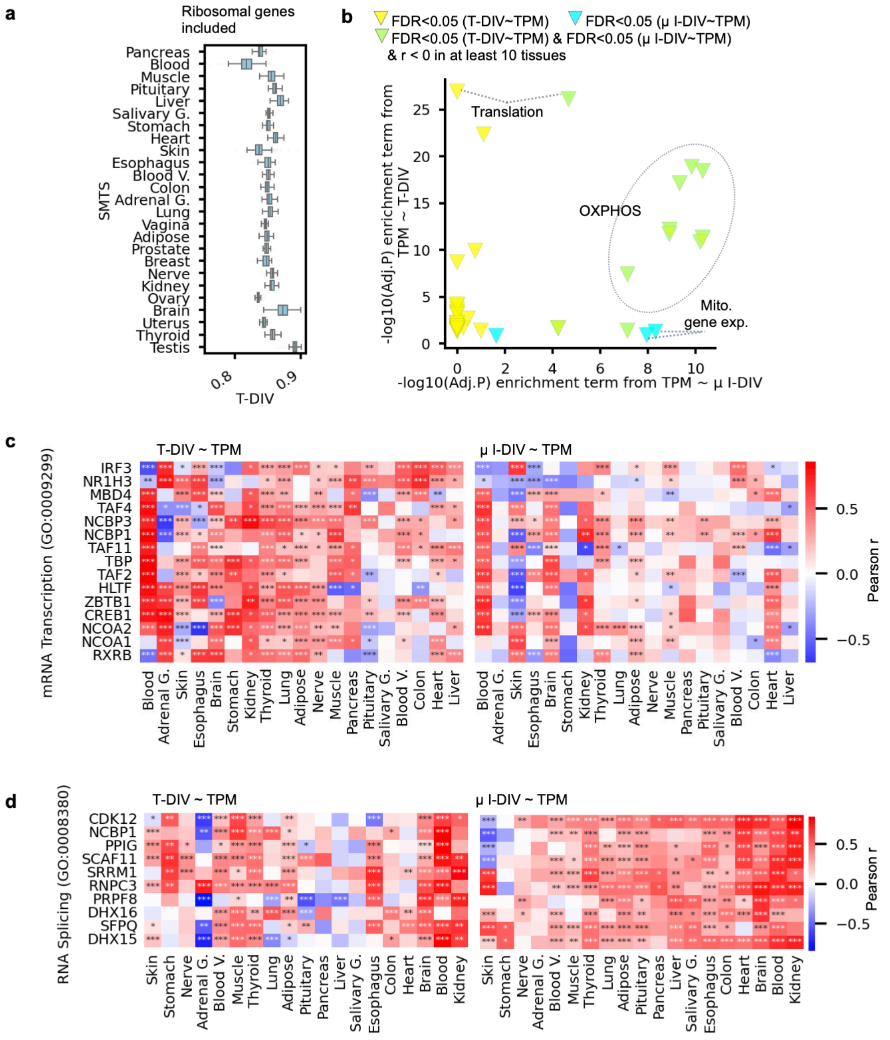


Supplementary Figure 4: Identification of genes whose expression is significantly and robustly associated to T-DIV or μ I-DIV.

(a) T-DIV calculated over expression of ribosomal genes, across 25 SMTS tissues. Box plots show the median (central line) and interquartile range (box, $25^{th}$–$75^{th}$ percentiles), with whiskers extending to the most extreme data point within 1.5x the interquartile range. One-way ANOVA $p<10^{-323}$. (b) Enrichr adjusted p-values of terms resulting from negatively correlating genes with μ I-DIV and/or T-DIV. -log10(FDR) of the enrichment of GO terms among genes correlated with T-DIV (y axis) are plotted against -log10(FDR) of the enrichment of GO terms among genes correlated with μ I-DIV (x axis). Each point represents one significantly enriched term (Enrichr FDR < 0.05) in at least one of the axes. Blue points represent terms significantly enriched among genes correlating with μ I-DIV only; yellow points represent terms significantly enriched among genes correlating with T-DIV only; green points represent terms significantly enriched among genes correlating with both metrics. (c) Heatmap shows Pearson correlation coefficients per gene to T-DIV (left) and to μ I-DIV (right), per gene and SMTS tissue. For the genes included in the enriched term mRNA transcription (GO:0009299; n = 15 genes). Asterisks indicate Pearson's p-values corrected for multiple testing (Benjamini–Hochberg): *** FDR < 0.001; ** FDR < 0.01; * FDR < 0.05; no asterisk, not significant (FDR ≥ 0.05). (d) Heatmap shows Pearson correlation coefficients per gene to T-DIV (left) and to μ I-DIV (right), per gene and SMTS tissue. For the genes included in the enriched term RNA splicing (GO:0008380; n = 10 genes). Asterisks indicate Pearson's p-values corrected for multiple testing (Benjamini–Hochberg): *** FDR < 0.001; ** FDR < 0.01; * FDR < 0.05; no asterisk, not significant (FDR ≥ 0.05).

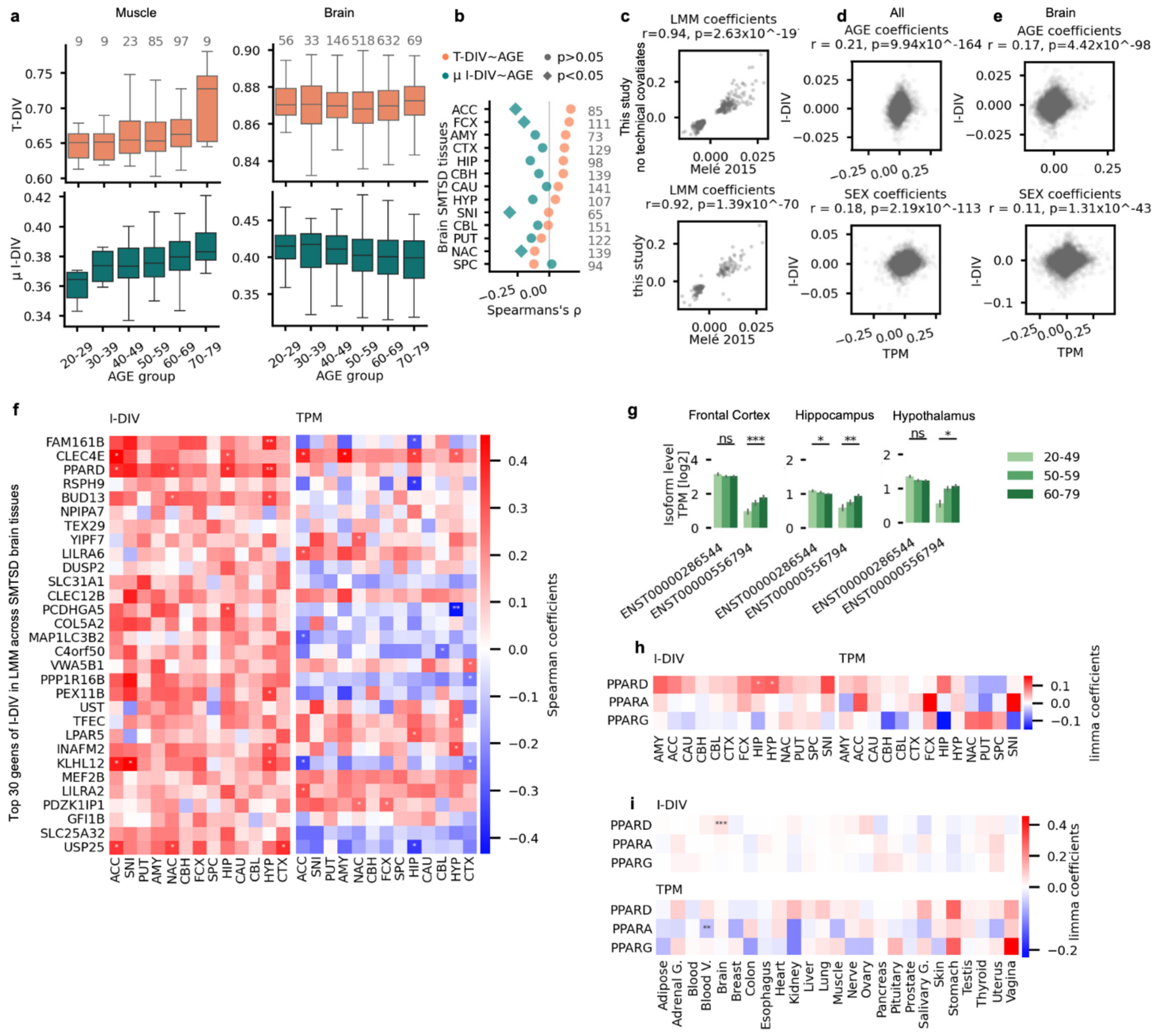


Supplementary Figure 5: Change in isoform diversity and gene expression with age.

(a) Box plots of the distribution of T-DIV (top) and μ I-DIV (bottom) in different age groups for muscle (left) and brain (right), showing the median (central line) and interquartile range (box, $25^{th}$–$75^{th}$ percentiles), with whiskers extending to the most extreme data point within 1.5x of the interquartile range. Number of samples per age group is stated above each box. (b) Spearman's correlation coefficients of μ I-DIV (green) or T-DIV (orange) with age for brain SMTSD tissue. Diamonds indicate significant coefficients (two-sided p-value < 0.05) and circles indicate non-significant coefficients (two-sided p-value ≥ 0.05). The number of samples per tissue is stated in grey. (c) Age coefficients of linear mixed models from current study plotted against age coefficients from Melé et al. Significant age coefficients in both studies are plotted (FDR-adjusted p-values by Benjamini–Hochberg < 0.05). Top: coefficients of current study excluding technical covariates (n = 424). Bottom: coefficients of current study including technical covariates (n = 176. Both are plotted against age coefficients from mixed linear model by Melé et al., 2015. Pearson's coefficients and two-sided p-values are stated above. (d)

Comparison of age and sex coefficients of linear mixed models of I-DIV (top) against TPM across all tissues. Top: Age coefficients comparison; Bottom: SEX coefficients comparison. Pearson's coefficients and two-sided p-values are stated above. (n=15834). (e) Comparison of age and sex coefficients of linear mixed models of I-DIV (top) against TPM across brain SMTSD tissues. Top: Age coefficients comparison; Bottom: SEX coefficients comparison. Pearson's coefficients and two-sided p-values are stated above. (n=15834). (f) Heatmaps show the age Spearman's coefficients per age and brain SMTS tissue I-DIV (left) and gene expression log(TPM+1) (right) for the top 30 genes ranked by I-DIV age coefficient which showed significance in I-DIV but not in TPM (FDR < 0.05) from the linear mixed model applied to brain regions. Asterisks indicate significant effects (Benjamini–Hochberg): ** FDR < 0.01; * FDR < 0.05. Brain SMTSD tissues: ACC, anterior cingulate cortex (BA24); AMY, amygdala; CAU, caudate (basal ganglia); CBH, cerebellar hemisphere; CBL, cerebellum; CTX, cortex; FCX, frontal cortex (BA9); HIP, hippocampus; HYP, hypothalamus; NAC, nucleus accumbens (basal ganglia); PUT, putamen (basal ganglia); SNI, substantia nigra; SPC, spinal cord (cervical c-1). (g) Bar plot of mean expression of each *FAM161B* isoforms per age group in Frontal cortex, hippocampus and hypothalamus (left to right). Error bars show the standard error; age groups are colored in shades of green from bright to darks. Significance of isoform changes measured in p-values from multiple linear regression (Methods): *** p < 0.001; ** p < 0.01; * p < 0.05; ns, not significant (p ≥ 0.05). (h) Heatmaps show the age coefficients from multiple linear regression per gene and brain SMTSD tissue in I-DIV (left) and gene expression log(TPM+1) (right) for *PPARD*, *PPARA* and *PPARG* genes. Asterisks indicate significant effects (Benjamini–Hochberg): * FDR < 0.05. Brain SMTSD tissues: ACC, anterior cingulate cortex (BA24); AMY, amygdala; CAU, caudate (basal ganglia); CBH, cerebellar hemisphere; CBL, cerebellum; CTX, cortex; FCX, frontal cortex (BA9); HIP, hippocampus; HYP, hypothalamus; NAC, nucleus accumbens (basal ganglia); PUT, putamen (basal ganglia); SNI, substantia nigra; SPC, spinal cord (cervical c-1). (i) Heatmaps show the age coefficients from multiple linear regression per gene and brain SMTS tissues in I-DIV (top) and gene expression log(TPM+1) (bottom) for *PPARD*, *PPARA* and *PPARG* genes. Asterisks indicate significant effects (Benjamini–Hochberg): *** FDR < 0.001.

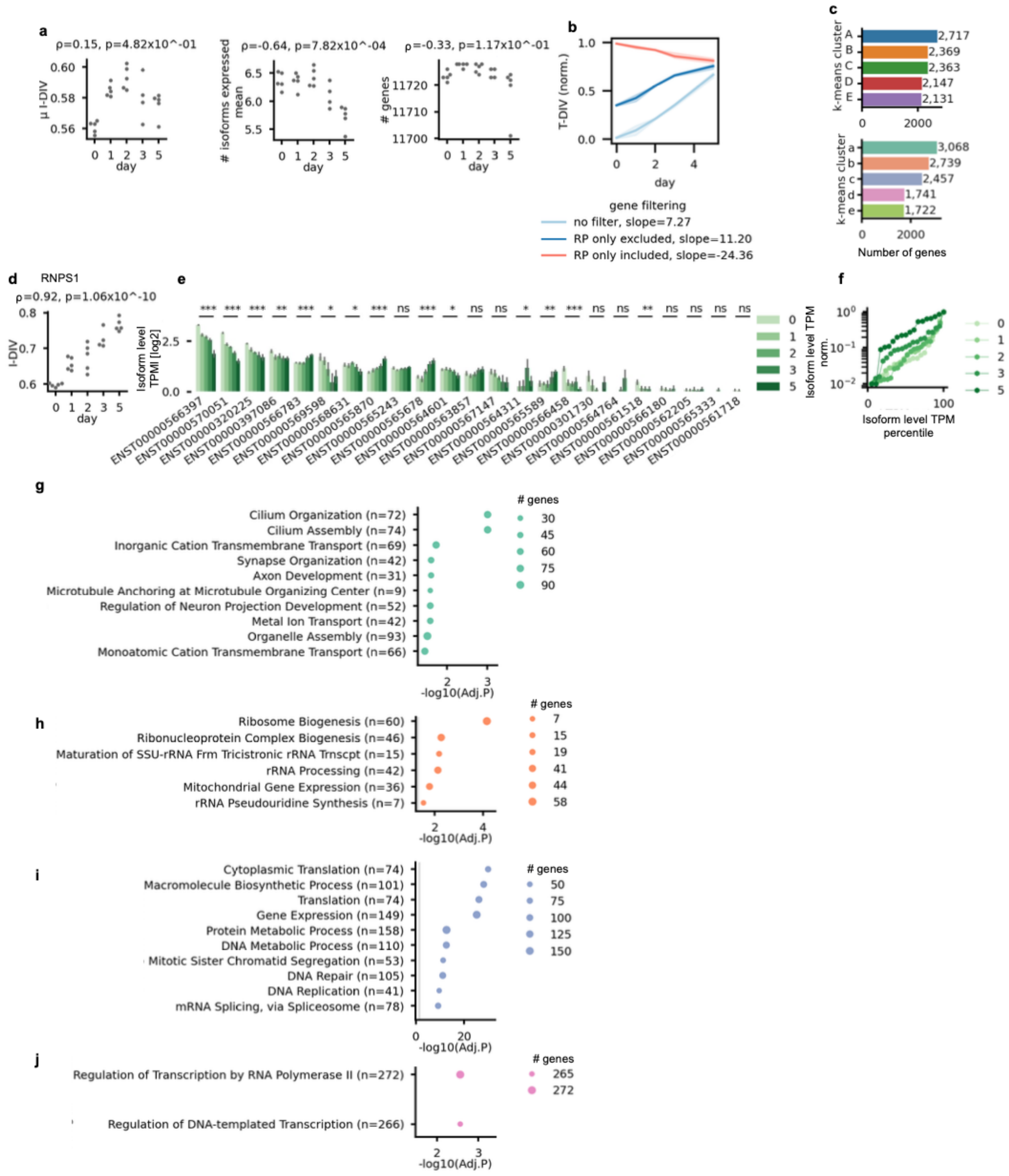


Supplementary Figure 6: Additional analyses of neuronal differentiation.

(a) Mean I-DIV, mean number of expressed isoforms and gene count shown by day of differentiation. Spearman's coefficient and two-sided p-value are stated above. (b) T-DIV calculated by three age groups: ribosomal protein genes (salmon), ribosomal protein excluded (dark blue), and all genes included (light blue) plotted against days of differentiation. (c) Bar plot of the number of genes k-means custer clustering based on I-DIV (top) and on gene expression (bottom). Colors correspond to the clusters in Fig. 5 panel d. (d) Swarmplot of *RNPS1* I-DIV at different days of differentiation. Number of samples per day, in x-axis order: 5, 5, 5, 4 and 5; Spearman's correlation coefficient and two-sided p-value is stated above. (e) Bar plot of mean expression of each *RNPS1* isoform per day of differentiation. Error bars show the standard error; days are colored in shades of green. Significance of isoform changes measured in p-values from multiple linear regression (Methods): *** $p < 0.001$; ** $p < 0.01$; * $p < 0.05$; ns, not significant. (f) Line plot of the scaled mean expression of the ten *RNPS1* isoforms, ordered by percentile of isoform expression across samples, for each day of differentiation (n=5, 5, 5, 4 and 5); days are colored in shades of green. (g-j) Enrichment analysis using GSEApy enrichr of each TPM cluster (n = 2,717, 3,068 2,739 , 2,457 and 1,741 genes in clusters a to d) against a background of all genes in the analysis (n = 11,727), showing -log10(FDR) for the top 10 most highly enriched GO terms across all clusters. Colors correspond to the k-means cluster bars in figure 5 panel d. The number of genes per term is indicated by size.
